# Crystal structures of alphavirus nonstructural protein 4 (nsP4) reveal an intrinsically dynamic RNA-dependent RNA polymerase fold

**DOI:** 10.1101/2021.05.27.445971

**Authors:** Yaw Bia Tan, Laura Sandra Lello, Xin Liu, Yee-Song Law, Congbao Kang, Julien Lescar, Jie Zheng, Andres Merits, Dahai Luo

## Abstract

Alphaviruses such as Ross River virus (RRV), chikungunya virus, Sindbis virus (SINV), and Venezuelan equine encephalitis virus are mosquito-borne pathogens that can cause arthritis or encephalitis diseases. Nonstructural protein 4 (nsP4) of alphaviruses possesses RNA-dependent RNA polymerase (RdRp) activity essential for viral RNA replication. No 3D structure has been available for nsP4 of any alphaviruses despite its importance for understanding alphaviral RNA replication and for the design of antiviral drugs. Here, we report crystal structures of the RdRp domain of nsP4 from both RRV and SINV determined at resolutions of 2.6 and 1.9 Å. The structure of the alphavirus RdRp domain appears most closely related to RdRps from pestiviruses, noroviruses, and picornaviruses. Hydrogen-deuterium exchange mass spectrometry (HDX-MS) and nuclear magnetic resonance (NMR) methods, showed that in solution, nsP4 is highly dynamic with an intrinsically disordered N-terminal domain. Both full-length nsP4 and the RdRp domain were capable to catalyze RNA polymerization. Structure-guided mutagenesis using a *trans*-replicase system identified nsP4 regions critical for viral RNA replication.

**Key Points:**

- Crystal structures of alphavirus nsP4 RNA polymerase domain from RRV and SINV.
- nsP4 protein is highly dynamic with an intrinsically disordered N-terminal domain.
- Optimized RNA elongation activity assay to facilitate antiviral discovery.

## Introduction

Alphaviruses (family *Togaviridae*) and flaviviruses (family *Flaviviridae*) comprise numerous important arthropod-borne viruses (arboviruses) that cause diseases in humans. Fast urbanization, massive international travel, and global warming have triggered the re-emergence of several arboviral diseases, such as dengue, Zika, and chikungunya fevers, affecting millions worldwide (1–3). Many arboviruses infect humans through *Aedes* mosquitoes which are abundantly present in tropical/subtropical regions and are expanding into temperate climate territories. Arbovirus members of the genus *Alphavirus* have been divided into Old World and New World alphaviruses, and cause clinically distinct symptoms (4,5). New World alphaviruses such as eastern equine encephalitis virus (EEEV) and Venezuelan equine encephalitis virus (VEEV) are primarily associated with encephalitis in horses and humans that causes death or long-term neurological impairment. Despite the generally nonfatal nature of infections, Old World alphaviruses, such as Chikungunya virus (CHIKV), cause a debilitating illness with persisting painful arthritis (6). Between March 2005 and April 2006, a major outbreak in the Reunion resulted in approximately 255,000 cases of chikungunya fever, approximately one-third of the total population of this French island. Human diseases caused by Ross River virus (RRV) and Sindbis virus (SINV) are generally milder, although they are also arthritogenic. Without any approved treatment, re-emerging outbreaks of alphaviruses highlight the unmet needs for vaccines and specific antiviral drugs. Available treatments for alphaviral infection are mostly symptom relief to minimize fever, joint pain, and associated inflammation. Several compounds, such as β-d-N 4-hydroxycytidine (NHC), favipiravir, and Compound-A, were claimed to be effective in inhibiting VEEV, CHIKV, and SINV RNA synthesis but have thus far not progressed towards clinical trials (7).

Alphaviruses have a positive-sense (+) RNA genome of approximately 11.8 kilobases in length with a 5’ type-0 cap and a 3’ polyadenylated tail. The two open reading frames (ORFs) in alphavirus RNA are flanked by *cis*-acting sequence elements: 5’ and 3’ untranslated regions (UTRs) and subgenomic (SG) promoters (8). RRV and CHIKV invade host cells by binding to the receptor Mxra8 (9), while SINV recognizes NRAMP as the main receptor (10). Receptor binding is followed by clathrin-assisted endocytosis, entry by fusion between the host cell and the viral membrane, and the release of the nucleocapsid containing the viral RNA genome into the cytoplasm. Upon its release, the genome functions as mRNA for polyproteins P123 and P1234 translated from its first ORF, and the expression of P1234 occurs through the readthrough of an inframe opal codon. These polyproteins are precursors for four nonstructural proteins (nsP1-4) that are formed via tightly regulated autoproteolysis and assemble into a viral RNA replication complex (RC). Alphavirus RCs are vesicular organelles called spherules and represent sites for viral RNA synthesis that are protected from the host immune response.

Three types of viral RNAs are synthesized during the infectious cycle: the full-length negative-strand intermediate, the full-length positive-strand genomic RNA, and the shorter positive-strand SG RNA. SG RNA synthesis is driven by the SG promoter, and the RNA is used to express structural proteins encoded by the second ORF of the alphavirus genome. The newly-made RNA genome and structural proteins assemble into progeny virions that are released by budding from the plasma membrane and are thereafter capable of further infection.

Alphavirus P1234 and its mature products contain all virus-encoded components required for RC assembly and all enzymatic activities needed for the synthesis and modification of viral RNAs. nsP4 contains RNA-dependent RNA polymerase (RdRp) activity responsible for *de novo* RNA synthesis and terminal adenylyltransferase (TATase) activity (11–13). In addition to nsP4, all other nsPs are also important for effective genome replication (12,13). Many molecular and structural details of the alphaviral nsP1-3 proteins have already been reported except for the C-terminal hypervariable region of nsP3, which is intrinsically disordered. In contrast, structural information for nsP4 has been lacking, in large part due to its intrinsic flexibility, which is important for its biological functions (14–22). Out of a total of ~610 amino acid residues, the N-terminal region of nsP4 (~ aa 1-100) comprises an N-terminal domain (NTD) unique to alphaviruses with a poorly defined function. The C-terminal region of the protein comprises the alphaviral RdRp domain. Over the years, several attempts to express nsP4 from alphaviruses in an active form for structural studies have yielded poorly soluble recombinant proteins with a tendency to aggregate. Substantial differences were noted, however, between nsP4 proteins of various alphaviruses: recombinant nsP4 of SINV possesses significant RNA polymerase activity (23), while the recombinant nsP4 RdRp domain of CHIKV displays only low template elongation activity (11). As a result, alphavirus nsP4 has resisted structural determination for several decades, which has hampered rational design for the development of compounds to target its activities. The availability of experimentally determined structures of nsP4 from any alphavirus would therefore constitute an important step to accelerate the high-throughput screening of inhibitors *in silico*, while the development of robust enzymatic RdRp assays would facilitate testing of these compounds. Furthermore, structural information on nsP4 would also be crucial for a better understanding of the structure and function of the alphavirus RC as a whole, and thus represent a strong foundation for further drug discovery efforts.

Here, we report crystal structures of alphavirus RdRp from both RRV and SINV determined at 2.6 and 1.9 Å resolution respectively (**Table 1**). Having a dynamic structure, the RRV RdRp contains several disordered segments and some electron density in the RNA binding tunnel remains unassigned. The SINV RdRp adopts an overall fold similar to RRV RdRp but appears more ordered: residues that were disordered in the RRV nsP4 crystal structure are now well resolved in SINV RdRp and almost fully occupy the central RNA binding tunnel. Thus, this alphavirus-unique RdRp conformation seems to be incompatible with RNA binding and polymerization. Complementary structural analysis performed using hydrogen-deuterium exchange mass spectrometry (HDX-MS) and nuclear magnetic resonance (NMR) helped to better understand nsP4 structural dynamics and revealed the intrinsically flexible conformations adopted by its NTD and RdRp domains in solution. Both full-length nsP4 (nsP4FL) and its RdRp domain displayed weak RNA polymerase activities *in vitro*. Finally, structure-guided mutagenesis experiments were used to gain functional insights into nsP4 in alphaviral RNA replication.

**Table 1.** Data collection, phasing and refinement statistics.

|  | RRV SeM-RdRp<sup>SER3</sup> | RRV RdRp<sup>SER3</sup> | SINV RdRp | SINV RdRp<sup>CS</sup> |
| --- | --- | --- | --- | --- |
| **Data Collection** | Se-SAD phasing | Native | Native | Native |
| **PDB ID** | - | 7F0S | 7VB4 | 7VW5 |
| **Space group** | P 2<sub>1</sub> 2<sub>1</sub> 2<sub>1</sub> | P 2<sub>1</sub> 2<sub>1</sub> 2<sub>1</sub> | P 1 | P 1 |
| **Unit cell dimensions** |  |  |  |  |
| a, b, c (Å) | 64.15 68.58 100.98 | 64.84 68.35 99.65 | 67 68.29 71.07 | 67.45 68.94 71.42 |
| α, β, γ (°) | 90 90 90 | 90 90 90 | 115.97 106.71 95.90 | 116.311 107.05 95.10 |
| **Wavelength** | 0.979 (peak) | 0.999 (native) | 1.000 | 0.946 |
| **Resolution range (Å)** | 46.85 - 2.83 (2.93 - 2.83) | 23.41 - 2.6 (2.69 - 2.6) | 40.52 - 1.9 (1.968 - 1.9) | 34.48 - 2.3 (2.382 - 2.3) |
| **Total reflections** | 296954 (29212) | 148510 (15410) | 524533 (36101) | 161646 (17056) |
| **Unique reflections** | 11129 (1107) | 14118 (1399) | 78572 (6646) | 45591 (4603) |
| **Mean I/sigma(I)** | 17.19 (0.30) | 23.81 (1.78) | 18.84 (1.90) | 13.25 (3.46) |
| **Completeness (%)** | 99.93 (99.82) | 99.66 (100.00) | 92.26 (71.47) | 94.41 (93.72) |
| **Multiplicity** | 26.7 (26.3) | 10.5 (11.0) | 6.7 (5.4) | 3.5 (3.7) |
| **R<sub>merge</sub><sup>b</sup>** | - | 0.066 (1.469) | 0.06941 (0.9519) | 0.05755 (0.3683) |
| **R<sub>meas</sub><sup>c</sup>** | - | 0.070 (1.544) | 0.07531 (1.054) | 0.06833 (0.4318) |
| **R<sub>pim</sub><sup>d</sup>** | - | 0.023 (0.470) | 0.02882 (0.4429) | 0.03631 (0.2233) |
| **CC<sub>1/2</sub><sup>e</sup>** | - | 0.984 (0.632) | 0.999 (0.708) | 0.998 (0.901) |
| **FOM** | 0.70 (0.66) | - | - | - |
| **Phase Error (°)** | 35.60 (40.21) | - | - | - |
| **Refinement Statistics** |  |  |  |  |
| **Reflections used in refinement** | - | 14112 (1399) | 78574 (5923) | 45652 (4415) |
| **Reflections used for R-free** | - | 696 (67) | 1858 (146) | 1169 (112) |
| **R<sub>work</sub><sup>f</sup>** | - | 0.265 (0.3877) | 0.1875 (0.3210) | 0.1950 (0.2564) |
| **R<sub>free</sub><sup>g</sup>** | - | 0.288 (0.4466) | 0.2239 (0.3549) | 0.2446 (0.3374) |
| **Number of nonhydrogen atoms** | - | 3109 | 8064 | 7648 |
| **macromolecules** | - | 3108 | 7160 | 7267 |
| **ligands** | - | - | 28 | 4 |
| **solvent (water)** | - | 1 | 876 | 377 |
| **Protein residues** | - | 419 | 917 | 930 |
| **RMSD (bonds)** | - | 0.006 | 0.003 | 0.002 |
| **RMSD (angles)** | - | 0.99 | 0.58 | 0.53 |
| **Ramachandran Statistics** |  |  |  |  |
| Ramachandran favored (%) | - | 91.81 | 97.45 | 97.6 |
| Ramachandran allowed (%) | - | 8.19 | 2.44 | 2.4 |
| Ramachandran outliers (%) | - | 0.00 | 0.11 | 0 |
| Rotamer Outliers (%) | - | 0.00 | 0.13 | 0.25 |
| Clashscore | - | 12.18 | 3.21 | 4.69 |
| Average B-factor | - | 108.36 | 28.32 | 44.61 |
| macromolecules | - | 108.37 | 27.71 | 44.54 |
| ligands | - | - | 34.43 | 52.67 |
| solvent | - | 69.40 | 33.15 | 45.86 |
| Number of TLS groups | - | 6 | 7 | 7 |

a. The values presented in parentheses are of the highest resolution shell.
where $R_{meas}$ is the precision indicator of individual observation for unmerged data.
where $R_{pim}$ is the precision indicator for the merged data
e. Correlation Coefficient, $CC =$

## Materials and Methods

### Molecular cloning

The sequences encoding RRV nsP4FL (aa 1-611) and its RdRp domain (aa 110-611) were obtained using PCR on an RRV-T48 cDNA plasmid template. The SINV nsP4FL (aa 1-610) and its RdRp domain (aa 91-610) were obtained using PCR on a SINV cDNA plasmid template (pSIN ReP5-GFP-GluR1 from Addgene). The PCR fragments were subcloned into the pSUMO-LIC vector using a ligation-independent cloning method in which the expressed recombinant protein had an N-terminal hexahistidine-linked small ubiquitin-like modifier protein (His_6_-SUMO) fusion tag which can act as solubility enhancer. For RRV RdRp, the RdRp surface entropy reduction construct (RdRp^SER3^) harboring two glutamine-to-alanine substitutions (Q192A and Q196A), nsP4FL polymerase-weakened mutant (GDD residues mutated to GNN; named as GNN) and construct for expression of NTD (aa 1-109) were obtained using site-directed mutagenesis (SDM) with the Toyobo KOD-FX Neo cloning kit. For SINV RdRP, the C164S mutant (named as SINV RdRp^CS^) was obtained via the same SDM method. These plasmids were propagated using the *E. coli* TOP10 strain.

Human RNA polymerase I (HSPolI) promoter-based plasmids for the production of replication-competent RNA templates of RRV (HSPolI-FG-RRV) and SINV (HSPolI-FG-SINV) have been previously described (24). Plasmids encoding P123 of RRV (CMV-P123-RRV) and SINV (CMV-P123-SINV) as well as plasmids expressing the corresponding nsP4 fused to ubiquitin at their N-termini (CMV-ubi-nsP4-RRV and CMV-ubi-nsP4-SINV) will be described elsewhere (25). Variants of CMV-ubi-nsP4-RRV and CMV-ubi-nsP4-SINV encoding the polymerase negative variant of nsP4 with GDD to GAA mutation in the polymerase active site were obtained using site-directed mutagenesis and designated CMV-ubi-nsP4^GAA^-RRV and CMV-ubi-nsP4^GAA^-SINV, respectively. Variants of CMV-ubi-nsP4-RRV and CMV-ubi-nsP4-SINV containing mutations listed in supplementary figure S6 were constructed using synthetic DNA fragments (Genscript). Sequences of all plasmids were verified using Sanger sequencing and are available from the authors upon request.

### Recombinant protein expression and purification

The recombinant proteins were expressed using the phage-resistant *E. coli* Rosetta II T1R strain developed in-house. Bacteria were grown in 2-4 L Luria-Bertani (LB) Millar Broth (Axil Scientific) supplemented with 50 μg/mL kanamycin (Gold Bio) and 34 μg/mL chloramphenicol (Gold Biotechnology) in a 37°C shaking incubator (INFORS) until reaching an optical density (OD_600_) measurement between 0.8 and 1.0. The shaker’s temperature was then adjusted to 18°C, expression was induced by the addition of isopropyl β-D-1-thiogalactopyranoside (IPTG) to a final concentration of 0.5 mM, and expression was carried out for 16-24 h. For selenomethionine labeling, an adapted protocol from Molecular Dimension was used for extended-expression at 18°C for 16 h (26). All the expression cultures were harvested by centrifugation for 10 min at 5000 x g, and bacterial pellets were frozen and stored at −80°C until recombinant protein purification. For ^13^C/^15^N-labeled NTDs from RRV, expression and induction were performed as described above except for the use of M9 minimal medium supplemented with 2 g/L ^13^C-glucose and/or 1 g/L ^15^NH_4_Cl.

The purifications of the recombinant nsP4 proteins (labeled or unlabeled) were conducted at 4°C using gentle treatments to ensure the structural integrity of the obtained samples. The bacterial pellet was suspended in IMAC A buffer (2X PBS, 500 mM NaCl, 5 mM β-mercaptoethanol, and 10% glycerol) supplemented with 1X protease inhibitor cocktail (DMSO free, MedChemExpress), 0.1-0.25% Tween-20 (Bio Basic) and 5 mM imidazole (Sigma Aldrich). The suspension was briefly sonicated before homogenization through 1000-bar pressurized PANDAPLUS2000 (GEA). The crude lysate was clarified via centrifugation in an ultracentrifuge XPN-100 (Beckman Coulter) at 235,000 x g. The clarified lysate was incubated with Ni-NTA FF beads (Bio Basic). Beads with bound recombinant protein were transferred to a column and subsequently washed with 15 column volumes (CVs) of IMAC A2 buffer (2X PBS, 500 mM NaCl, 5 mM β-mercaptoethanol, 10% glycerol, and 10 mM imidazole). The recombinant protein was eluted in 5 CV of IMAC B buffer (2 X PBS, 500 mM NaCl, 5 mM β-mercaptoethanol, 10% glycerol and 300 mM imidazole) and treated with Ulp1-SUMO-protease (1:25 w/w ratio of protease: protein) for 16-20 h at 4°C in the presence of 250 mM NaCl to remove the N-terminal His_6_-SUMO tag. The cleaved recombinant protein was loaded onto a Heparin 5 HP column (Cytiva) at an adjusted concentration of 100 mM NaCl and subsequently eluted using 900 mM NaCl. The eluate was further separated through size-exclusion chromatography (SEC) in a HiLoad S75 or S200 16/600 Superdex^®^ column (Cytiva) on an AKTA Pure 25 M fast protein liquid chromatography (FPLC) system (Cytiva).

Products obtained at all purification steps were examined for their purity by SDS-PAGE. The fractions corresponding to the elution peak were selected and concentrated to 2-50 mg/mL through the use of Amicon^®^ Ultra-15 Centrifugal Unit (Merck), frozen and stored at −80°C for downstream experiments. The storage buffer is 25 mM HEPES pH7.4, 300 mM NaCl, 2 mM tris(2-carboxyethyl)phosphine (TCEP), 5% Glycerol.

### X-ray crystallography

Crystallization conditions for purified recombinant nsP4 proteins were screened with Hampton Research commercial kits (PEG-ION HT, PEG-RX HT, INDEX HT, SALT-RX, CRYSTALSCREEN HT) at concentrations 100-121 μM with and without ligands, aided by a Mosquito robotic liquid handler (TTP Lab) and crystallization drops were periodically imaged in the RockMaker (Formulatrix^®^) maintained at 20°C.

Under optimized conditions, 121 μM RRV RdRp was incubated with 2-5 mM MgCl_2_, 2-5 mM deoxyadenosine triphosphate or adenosine triphosphate (dATP/ATP) and 135 μM synthesized self-annealing duplex-RNA R2 (dsRNA: 5’ GCAUGGGCGCCC 3’) for 1-2 h. This protein-RNA mixture led to crystal formation with sizes of 0.2-0.5 mm on a vapor-diffusion hanging-drop of 24% (w/v) PEG 3350, 0.1 M Na HEPES/Tris HCl pH 8.0, 0.2 M ammonium acetate, and 3% trimethylamine N-oxide (TMAO) in 5-7 days. The crystals were dehydrated using the same buffer supplemented with 30% (w/v) PEG 3350 and 10% glycerol as cryoprotectants before snap freezing on a cryoloop in liquid nitrogen. Crystals of SINV RdRp (both wild type and C164S mutant) were obtained with a concentration of 110 μM incubated with 2 mM ATP, 2 mM MgCl_2_, 18-20% PEG3350 and 0.2 M magnesium acetate. These crystals grew to a final size of 0.3-0.5 mm under vapor-diffusion hanging drop and were snap-frozen with the addition of 10-20% glycerol.

X-ray diffraction datasets (**Table 1**) were collected at Taiwan Photon Source (TPS) Beamline TPS-05A, Japan Spring-8 Beamline BL32XU (27,28), Swiss Light Source Beamline X06DA (PXIII), and Australian Synchrotron Beamline MX2 (29,30), all with a nitrogen stream to maintain crystals at 100 K. Data were collected for at least a total consecutive sweep of 200 degrees with an oscillation step of 0.1-0.5 degrees and a 70-90% attenuated beam. The detector was placed at a distance of 260-300 mm from the crystal for native data collection at wavelengths of ~0.95-1 Å. Single-wavelength anomalous diffraction data collected at the selenium K-edge for the seleniated RRV RdRp protein were used for SAD phasing and the Semet positions were used as markers to aid model building. Data were preprocessed on-the-fly by an XDS-based program (31) and further processed in the *CCP4 7.1* (32) and *Phenix* 1.19 software suites (33) for SAD phasing and automated model building. Crystal structures were displayed at the computer graphics using *Coot* and refined with *Phenix* (33,34). Table 1 summarizes the crystallographic refinement parameters.

### Hydrogen-deuterium exchange detected by mass spectrometry

For both RRV and SINV, 5 μM recombinant of either nsP4FL or RdRp or NTD protein in a buffer containing 50 mM Na HEPES, pH 7.4, 150 mM NaCl, 5% glycerol, 5 mM MgCl2, and 2 mM DTT was used to carry out HDX reactions at 4°C. Four microliters of protein/protein complex with ligand/peptide were diluted in 16 μL D_2_O in exchange buffer (50 mM Na HEPES, pH 7.4, 50 mM NaCl, and 2 mM DTT), incubated for various HDX time points (e.g., 0, 10, 60, 300, 900 s) at 4°C and quenched by mixing with 20 μL of ice-cold 1 M TCEP 100 mM NaH_2_PO_4_. Each quenched sample was immediately injected into the LEAP Pal 3.0 HDX platform. Upon injection, samples were passed through an immobilized pepsin column (2 mm□× 2 cm) at 120 μL min^−1^ and the digested peptides were captured on a C18 PepMap300 trap column (Thermo Fisher) and desalted. Peptides were separated across a 2.1 m□×□5 cm C18 separating column (1.9 mm Hypersil Gold, Thermo Fisher) with a linear gradient of 4-40% CH_3_CN and 0.3% formic acid over 6 min. Sample handling, protein digestion, and peptide separation were conducted at 4°C. Mass spectrometric data were acquired using a Fusion Orbitrap mass spectrometer (Thermo Fisher) with a measured resolving power of 65,000 at m/z 400. HDX analyses were performed in duplicate or triplicate with single preparations of each protein state. The intensity-weighted mean m/z centroid value of each peptide envelope was calculated and subsequently converted into a percentage of deuterium incorporation. Statistical significance for the differential HDX data was determined by an unpaired Student’s t-test for each time point, a procedure that is integrated into the HDX Workbench software (35). Corrections for back exchange were made based on an estimated 70% deuterium recovery and accounting for the known 80% deuterium content of the deuterium exchange buffer.

For peptide identification, Product ion spectra were acquired in data-dependent mode with the top eight most abundant ions selected for the production analysis per scan event. The MS/MS data files were submitted to Proteome Discover 2.4 (Thermo Fisher) for high-confidence peptide identification. For data rendering, The HDX data from all overlapping peptides were consolidated to individual amino acid values using a residue averaging approach. Briefly, for each residue, the deuterium incorporation values and peptide lengths from all overlapping peptides were assembled. A weighting function was applied in which shorter peptides were weighted more heavily and longer peptides were weighted less. Each of the weighted deuterium incorporation values was then averaged to produce a single value for each amino acid. The initial two residues of each peptide, as well as prolines, were omitted from the calculations.

### NMR data collection and analysis

The NMR spectrum of the RRV NTD was collected at 298 K on a Bruker Avance II magnet with a proton frequency of 600 MHz equipped with a cryoprobe. A^13^C/^15^N-labeled (0.3 mM) NTD in a buffer that contained 20 mM Na-PO_4_, pH 6.5, 150 mM NaCl, and 1 mM DTT was used for data acquisition. Experiments including 3D-HNCACB, HNCOCACB, HNCA, HNCOCA, and HNCO were collected for resonance assignment. Assignment was challenging due to the absence of signals from several residues. Data were processed using NMRPipe (36) and visualized using NMRView (37).

### Structure analysis

The 3D crystal structures of RRV and SINV RdRp (this work) were queried against the full PDB database through the Dali server (Dali; http://ekhidna2.biocenter.helsinki.fi/dali/) for matching structural homologs. Amino acid sequences of the top hits from Dali were aligned through the COBALT Multiple Alignment Tool (https://www.ncbi.nlm.nih.gov/tools/cobalt/cobalt.cgi) or MUSCLE (https://www.ebi.ac.uk/Tools/msa/muscle/) to generate a multiple sequence alignment (MSA) file. The sequence-conservation map overlay with the RdRp secondary structure was generated through ESPRIPT (https://espript.ibcp.fr) (38) with the input of the MSA file and RdRp^SER3^ PDB. The nsP4FL, RdRp, and NTD homology models of RRV or SINV nsP4 were predicted based on comparative modeling (CM) mode with the template of the best-refined RdRp^SER3^ structure by sampling for 5 models through the Robetta server (https://robetta.bakerlab.org/) (39). The homology models were superimposed on other experimental crystal structures for comparison and analysis. These homology models were colored and annotated for the HDX-MS experiment. All structural displays were created in PyMOL 2.4.0 (Schrodinger LLC.) (40).

### RNA polymerase assay

The optimized RNA polymerase assay from (11) was conducted here using 1 μM recombinant nsP4 proteins and 50 nM internally labeled fluorescent T1 hairpin-RNA template (5’ FAM-UUUUUUUUUUAGGACCGCAUUGCACUCCGCGGUCCUAA 3’) in PA buffer containing 25 mM Na HEPES pH 8.0, 2.5 mM MnCl_2_, 5.0 mM MgCl_2_, 20 mM KCl, 5 mM TCEP, 3 mM LDAO, 0.01% Triton X-100, and 2 U/μL murine RNase inhibitor (NEB) in nuclease-free water (HyClone). Reactions were initiated with 2 mM ATP and incubated for 0 to 24 h at 25°C in the dark. At selected incubation time points, the reactions were quenched using 2X RNA loading dye (95% formamide, 0.02% SDS, 0.01% xylene cyanol, 0.02% bromophenol blue, and 1 mM EDTA). Samples were loaded onto 17.5% urea-PAGE and run at 200 V for 1.5-2 h. The urea-PAGE was imaged on a Bio-Rad Gel-Doc XR+ Imager using the Fluorescence Blot ALEXA488 protocol to analyze the RNA polymerase activity.

### *Trans*-replication assay

U2OS human bone osteosarcoma cells (ATCC HTB-96) were maintained in Iscove’s Modified Dulbecco’s *medium* (Gibco) containing 10% FBS, 2 mM L-glutamine, 100 U/mL penicillin, and 0.1 mg/mL streptomycin at 37°C in a 5% CO2 atmosphere. Cells grown on 12-well plates were cotransfected with the following combination of plasmids (each used at 1 μg per transfection): HSPolI-FG-RRV, CMV-P123-RRV, CMV-ubi-nsP4-RRV or its mutant versions (Supplementary Data, Fig. S6) for RRV *trans*-replicase and HSPolI-FG-SINV, CMV-P123-SINV, CMV-ubi-nsP4-SINV or its mutant versions for SINV *trans*-replicase. Transfections were performed using Lipofectamine LTX with PLUS reagent (Thermo Fisher Scientific) according to the manufacturer’s instructions. Transfected cells were incubated at 37°C for 18 h. All transfections were performed in triplicate, and experiments were repeated at least twice. After incubation, cells were lysed, and the activities of Fluc and Gluc were measured using the Dual-Luciferase-Reporter assay (Promega). Fluc and Gluc activities measured for cells transfected using plasmids expressing active replicases or their mutants under investigation were normalized to those obtained for the corresponding control cells transfected with HSPolI-FG-RRV, CMV-P123-RRV, and CMV-ubi-nsP4^GAA^-RRV or HSPolI-FG-SINV, CMV-P123-SINV and CMV-ubi-nsP4^GAA^-SINV.

## Results

### Determination of the crystal structures of nsP4 RdRp from RRV and SINV

We obtained soluble recombinant nsP4FL and RdRp proteins of both RRV and SINV (**Figure 1a**) using the *E. coli* expression system. The RdRp recombinant proteins, lacking the N-terminal 109 or 90 amino acids of nsP4 respectively, were purified as soluble active enzymes that could be crystallized. The size exclusion chromatography profiles of these proteins indicate they are monomeric in solution. Crystals of the RRV RdRp initially diffracted to a maximum resolution of 3.8-4.0 Å. To improve RRV RdRp crystals diffraction, we substituted hydrophilic glutamine residues (Q192 and Q196) that are presumably exposed at the surface of the folded RdRp protein with alanines. The obtained surface-entropy double mutant RdRp^SER3^ formed crystals with an increased size that allowed native X-ray diffraction intensities to be collected to a maximum resolution of 2.60 Å (**Table 1**). Because no convincing solution was found using molecular replacement (MR), L-selenomethionine (SeM)-labeled RRV RdRp^SER3^ was used for single-wavelength anomalous diffraction (SAD) phasing at the Se K edge. SAD data were collected at 2.83 Å resolution using a crystal isomorphous to native RdRp^SER3^ crystals (**Table 1**). A total of 419 out of 502 residues could be built, including eighteen α-helices, three 3_10_ helices, and ten β-strands (**Figures 1B-C and Table 1**). Of note, no bound ligands, such as RNA, ATP, Mg^2+^ or trimethylamine N-oxide (TMAO), were identified in the RdRp^SER3^ crystal structure despite their presence under crystallization conditions (see Methods). An anomalous Fourier difference map using data collected at the Se K-edge allowed unambiguous mapping of eleven Se-Met residues (out of a total of thirteen) onto the structure, giving confidence for the correctness of the atomic model (**Figure S1**). While we were making progress in building and refining the RRV RdRp crystal structure, we managed to obtain crystals of the wild-type SINV RdRp spanning amino acids 91-610 in a crystallization condition different from the RRV protein (see Methods). SINV RdRp crystals diffract to 1.9 Å resolution and belong to the P1 space group with two RdRp molecules per asymmetric unit (ASU) (**Table 1 and Figure S3**). The SINV RdRp structure was readily determined by molecular replacement using the RRV RdRp structure as a search probe and refined to a R_free_ value of 22.4% with good stereochemistry (**Table 1, Figures 1 and S3**).

**Figure 1.**
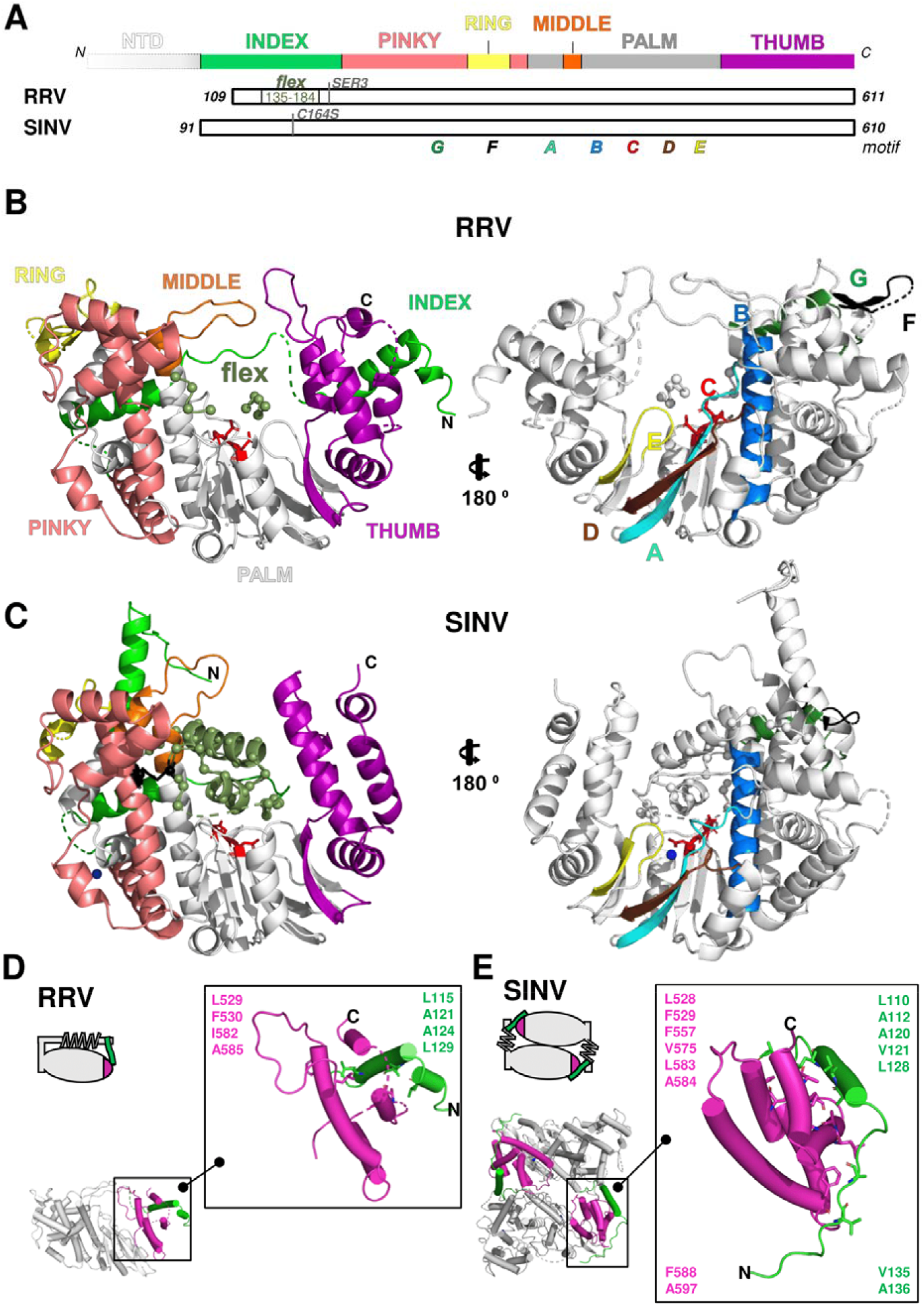
Crystal structure of Alphavirus nsP4 RdRp domain. (A) Schematic presentation of RRV and SINV nsP4. The crystallised regions of nsP4: SINV RdRp (aa 91-610) and RRV RdRp (aa 109-611) are annotated. Their flexible region (aa 135-184; denoted as *“flex”* in smudge green) occupying the central tunnel, and (sub)domain distribution is shown as follows: n-terminal domain (NTD) in white and RdRp domain consisting of fingers with colored fingertips (index in green, middle in orange, ring in yellow and pinky in pink), palm in gray and thumb in purple. The mutations targeting disulfide-bond forming cysteine (C164S) in SINV RdRp and surface-entropy site (labeled as SER3 to represent surface-entropy mutations of Q192A and Q196A) in RRV RdRp are marked. The locations of the A-G motifs are shown in colors (colored as A: teal; B: marine; C: red; D: brown; E: yellow; F: black; G: forest green). (B-C) The annotated crystal structures of RRV RdRp (RdRp^SER3^) in (B) and SINV RdRp (wild type) in (C) are colored according to (A) for displaying their subdomain spatial distribution from the front right-hand view (left panel) and their motifs in the 180° rear view (right panel). Motif C that comprises conserved GDD active-site residues is highlighted in red sticks. Polyalanine considered in the flex region of RRV RdRp^SER3^ and built residues in the SINV RdRp flex region are both represented as a smudge-green ribbon which is traced by backbone Cα residues as a sphere. Additional features in SINV RdRp in (C): the cysteines (C164 in *flex*, C323 in pinky) are labeled as black sticks (left panel) while magnesium cations are colored navy blue. (D-E) The contrast of RdRp crystallographic configurations for (D) monomeric RRV and (E) dimeric SINV is presented in both crystal structure (top view) and simplified block (non-hydrophobic-interacting part colored as gray and zig-zag part symbolizes *flex* region) to present the hydrophobic interactions between the index finger (green region) and thumb (magenta region) on the left panel respectively. The hydrophobic interface of RRV and SINV between the N-terminal index finger and C-terminal thumb region is highlighted in the respective zoom-in box on the right panel of both (D) and (E). In these zoom-in boxes, all hydrophobic residues (hydrophobic interaction distance within 4Å) are shown as sticks and listed accordingly to the color of the index finger and thumb. The top list is the common residues shared between RdRp of RRV and SINV while the bottom list is SINV-RdRp-unique residues at the hydrophobic interface.

### Structural comparison of the two alphavirus RdRps

Both RRV and SINV RdRps adopt the characteristic right-hand fold of viral RdRps, where fingers-palm-thumb subdomains are sequentially arranged from the amino-to-carboxylic ends. The overall structure of SINV RdRp is similar to RRV within an RMSD of 1.97 Å (417 Cα)(**Table 1, Figure 1 and S3**). The core catalytic domain from both RRV and SINV RdRps including the palm region is well defined in continuous electron density and structurally well conserved between both RdRp proteins following superposition (**Figure 1B-D and Table S1**). Compared to RRV RdRp, the higher-resolution structure of the SINV RdRp reveals more details of the structure and also bound solvent molecules (magnesium and glycerol). Two magnesium cation binding sites at aspartates of residue 392 and 466, in which the latter is part of the active site GDD motif (**Figure S3A**).

A total of 83 residues are missing from the highly dynamic RRV RdRp crystal structure, while there are only 39 missing residues from the SINV RdRp structure. The 83 mobile residues in RRV RdRp belong to the N-terminal fingertips and C-terminal thumb subdomains: index finger (residues 135-184; this flexible region is denoted as “flex” region hereafter) (**Figure 1D and S2C**), pinky, and ring fingers (residues 210-213 and 292-307, respectively) **(Figure S2A**), and C-terminus of the protein (residues 575-578, 588-593 and 600-611) (**Figure 2A and Figure S2B**). Conversely, missing residues in SINV RdRp are located in N-terminal fingertips (index residues 91-103, flex residues 167-178, pinky residues 206-210, and ring residues 288-307) and C-terminus residues (residues 603-611) **(Figure 2**). The relatively large number of missing residues of the RRV RdRp structure suggests a highly dynamic structure which is also detected in the HDX measurement (see below, **Figure 2**).

**Figure 2.**
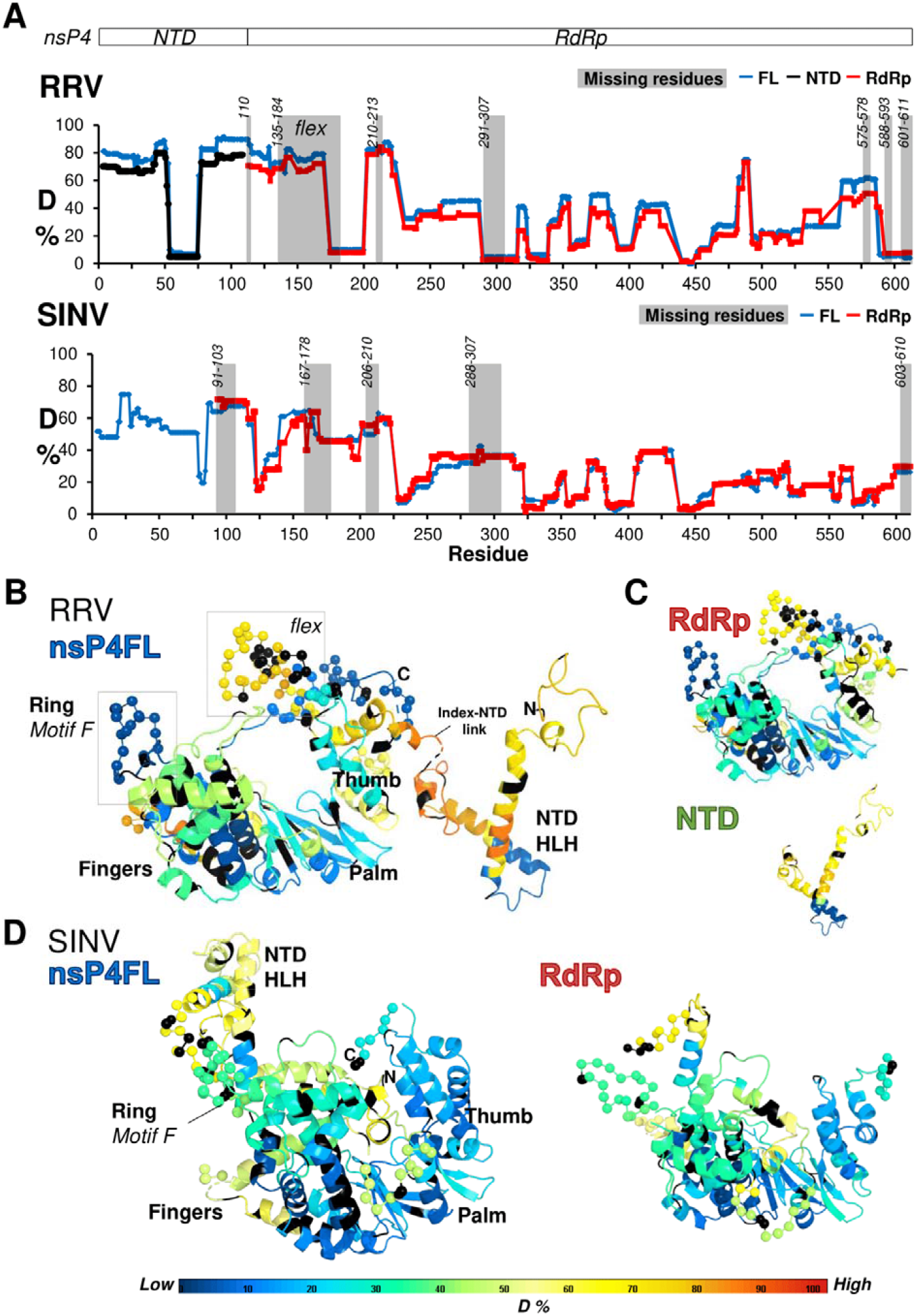
Structure dynamics of RRV nsP4 in solution. (A) The HDX profiles of the recombinant constructs of both RRV and SINV: nsP4FL (blue), RdRp (red), and NTD (green), in solution. On the graph, the deuteration incorporation levels expressed in percentage (D%) are plotted against the nsP4 residue numbers (RRV: aa 1-611, SINV: aa 1-610). The residues missing in the resolved structure are shaded as gray areas, and their positions are indicated. The longest unresolved region in RRV RdRp (residues 135-184) is designated as “flex”. (B-C) The color-coded ternary structures of Robetta homology models for RRV (B) nsP4FL, (C) RdRp and NTD, and for SINV nsP4FL and RdRp in (D), according to the representation of D% in rainbow heatmap to visualize structural flexibilities. These models were generated via Robetta (39) to fill the missing residues not covered in the current crystal structures, including the flexibly orientated NTD. The HDX-protection profiles of RRV nsP4 in (A) were color-mapped to these RRV homology models at single-amino-acid resolutions to guide the visual analyses and 3D comparisons between nsP4FL and RdRp and NTD. RRV nsP4FL is annotated with N/C-terminus, subdomains, motifs, and highlighted regions (Ring Motif F, Pinky, *flex*, Index-to-NTD linker, HLH substructure) as a reference to other models in (C) and (D). The missing residues are presented as Cα spheres in these homology models. The residues colored in black are those not covered/excluded by HDX data rendering. Rainbow bar as reference for (B),(C), and (D): D% is visualized in the rainbow heatmap spectrum, where the red end represents high D% (structured region) and the blue end represents low D% (disordered region).

There are noticeable structural differences in the fingers and thumb subdomains. The highly flexible fingers subdomain of RRV (subdivided into the index, pinky, ring, and middle finger subdomains) is significantly different from the corresponding SINV region with an RMSD of 1.90 Å (**Fig. 1B-D and Table S1**). The RRV RdRp fingertips extend from the fingers towards the thumb to form the distinctive encircled ring conformation for RNA polymerization first seen in the HCV polymerase structure (41,42) To fold into this topology, the RRV nsP4 index finger extends its N-terminal helix horizontally towards the thumb to which it docks via the formation of a hydrophobic interface (**Figure 1B-D and Figure S3B-D**). The flexible SINV index fingertip appears detached from the core domain and crosses over to a neighboring RdRp molecule within the ASU, forming a dimeric configuration within the crystal (**Figure 1B-D and Figure S3B-D**). This difference between RRV and SINV index fingers suggests that the N-terminus end is dynamic and potentially undergoes structural reorganization during RC assembly and RNA loading. However, this dimeric form of SINV nsP4 RdRp is likely a crystallographic artifact, as all reported alphaviral nsP4 proteins are monomeric in solution based on the present and past gelfiltration data (12,13,43).

While the Flexi region of the RRV RdRp is largely disordered, the equivalent region of SINV RdRp folds into the central RNA binding and catalysis tunnel (**Figure 1 and Figure S3**). Unexpectedly, a disulfide bond between cysteine residues C164 and C323 is formed. The cysteine-to-serine residue mutation (C164S) at the less conserved residue C164 did not alter the enzyme’s overall folding or catalytic ability so this disulfide bond appears not to be important for virus replication (**Figure 1C and S7C-D**). Other fingers such as middle, ring, and pinky fingers are more conserved structurally between the two RdRps. The middle finger of RRV RdRp maintains the classical RdRp helix-loop-helix substructure but with its loop orienting lower than that of SINV RdRp, in which the middle and index fingers interact (**Figure 1D**). The ring and pinky fingers are the most conserved fingertips between the two RdRps. Importantly, the occupation of the central tunnel by residues from the finger domain would exclude RNA substrates (template or primer) binding in both RRV and SINV RdRps. Therefore, nsP4 must undergo a very large conformational rearrangement of the finger domain to accommodate the RNA template and to catalyze viral RNA polymerization. Overall, the finger subdomains indicate flexibilities for conformational changes, likely for its enzymatic roles, nsPs interactions, and viral RNA binding.

Further, the thumb subdomain in the RdRps of SINV and RRV has limited structural similarity in their α-helical arrangement (**Figure 1D**). The loop between the first two α-helices (a1 and a2) is longer in RRV (residues 525-557) showing higher flexibilities than the equivalent of SINV with a shorter loop and more structured α-helices. The unbuilt regions in the C-terminus end of RRV RdRp (**Figure 1D**) due to weak electron densities also contributes to the increased RMSD despite the spatial outlines of the α-helices in both RdRps of RRV and SINV remains inframe. The higher dynamics in RRV RdRp thumb region observed here are again supported by comparing the HDX profiles of RRV and SINV nsP4 (see below, **Figure 2**).

In summary, the conformations captured in the present crystal structures are likely to have been partly induced by crystal packing forces leading to shifts in the finger and thumb domain compared to catalytically fully competent forms. Moreover, beside crystal packing forces, the structural context (eg interactions with other members of the alphavirus RC is also likely to be important to determine the RdRp conformation, given the high flexivbility of the isolated recombinant protein.

### The structural dynamics of nsP4 and its domains in solution

To gain insights into the conformational dynamics of nsP4 and to probe intramolecular interactions in solution, we performed HDX-MS and NMR studies on recombinant nsP4 and its domain proteins. HDX-MS is a proteomics approach to study protein dynamics by measuring the deuterium-hydrogen exchange rates between solvent D_2_O and the target protein. A high percentage of deuteration (D%; hydrogen exchanged to deuterium) reflects the high solvent accessibility and flexibility of the regions on RRV and SINV nsP4. To illustrate the structural dynamics in 3D, we plotted the HDX profiles in D% in heatmap format to these structural models for comparison (**Figures 2B-D and Figure S4-5**). The missing residues in the crystal structures were modeled using Robetta ternary-structure prediction (39). Based on the high D% HDX profile observed, the NTD of RRV nsP4FL appears quite flexible in solution, except at residues 53-74, where a predicted helix-loop-helix (HLH) substructure is located (**Figure 2B-C)**. The linker between NTD and RdRp appears to have higher D% values, indicating increased dynamics in solution (**Figure 2A**). Given its low molecular mass of 12.6 kDa, the recombinant NTD is amenable to structural characterization via NMR in solution. The NTD selected from RRV contains 107 amino acids with an expected number of 99 cross-peaks (excluding Pro residues) in the ^1^H-^15^N-HSQC spectrum. From these, approximately 60 cross-peaks could be observed, making backbone resonance assignments challenging (**Figure S6**). The narrow dispersion of the cross-peaks in the spectrum suggests that the NTD contains flexible and helical regions that support the NTD homology model and are in agreement with the HDX profile (**Figure 2A, 2C and S6**).

The palm and thumb subdomains as well as the middle and ring fingers are largely protected from the deuterations and therefore constitute a buried structural core. In contrast, the flexible N-terminal index and pinky fingers harboring the respective *flex* region and motif G (residues 215-221) and the thumb C-terminus region (residues 560-585) became highly deuterated in the HDX experiment (**Figures 2A-C and Figure S4**). Thus, these are less-protected regions, a finding that is in agreement with their high mobility found in the crystal structure of RRV RdRp (**Figure 1B**). The HDX protection profile of the ring fingers and the last 20 amino acids at the protein C-terminus is in contrast with the RRV RdRp crystal structure, where these regions had weak or missing electron densities.

Overall, the similar HDX profiles of the NTD protein by itself and within the nsP4 from both SINV and RRV indicate that there is no strong intramolecular interaction between the NTD and the RdRp region of the nsP4.

### Comparison of the RRV RdRp structure with that of other viral RdRps

The alphavirus nsP4 polymerase domain of RRV and SINV adopts a conserved tertiary structure harboring features common to viral RdRps. A structural homology search of the Protein Data Bank (PDB) database (44) revealed that the closest structural homologs of both RRV and SINV RdRps are RdRps from the human Norwalk virus from the *Caliciviridae* family (NV; PDB 5TSN) (45), enterovirus-71 (EV71, PDB 6KWQ) from the *Piconaviridae* family (46) and classical swine fever virus (CSFV, PDB 5Y6R) as well as bovine viral diarrhea virus (BVDV) from the *Flaviviridae* family (47) (**Figure 3**). In RRV nsP4, the conserved polymerase motifs forming a fully encircled right-hand fold are arranged in the primary structure as G-F-A-B-C-D-E, showing evolutionary conservation with the *Picornaviridae* (but not to the birnavirus) RNA polymerase (42) (**Figure 1 and Figure 3**). The SINV nsP4 did not form the encircled right-hand fold due to the index finger interacting with the neighbor crystallographic ASU but this intermolecular contact does not affect the overall structure and their motif arrangements. The core polymerase domain at the palm of alphaviral RdRp contains motifs A, B, C, and E that are structurally well-conserved compared to known positive-strand RdRps (**Figure 3**). The Phe residue on motif E of alphaviral nsP4 points towards the GDD residues in the β-turn of motif C, as in other viral RdRps (**Figure 3**; motif E in yellow and motif C in red). In contrast, motif D is likely to constitute an alphavirus distinctive feature that separates it from other viral RdRps. Residues from motif D of nsP4 adopt an elongated β-strand conformation, which forms a Greek-key-like β-sheet motif, next to motifs A-C (**Figure 3A**). The least conserved motifs lie in the pinky finger (motif G) and ring finger (motif F) subdomains which are known for poor sequence homology in viral RdRps (**Figure 3B**). The fingertips are typically folded in β-sheets extending towards the thumb like a lid over the palm. The thumb subdomain of nsP4 shows poor structural homology with RdRp homologs, and its C-terminal region is folded in a different orientation without β-sheet formation. These structural differences make the alphavirus nsP4 an α-helix rich structure compared to its closest viral RdRp homologs.

**Figure 3.**
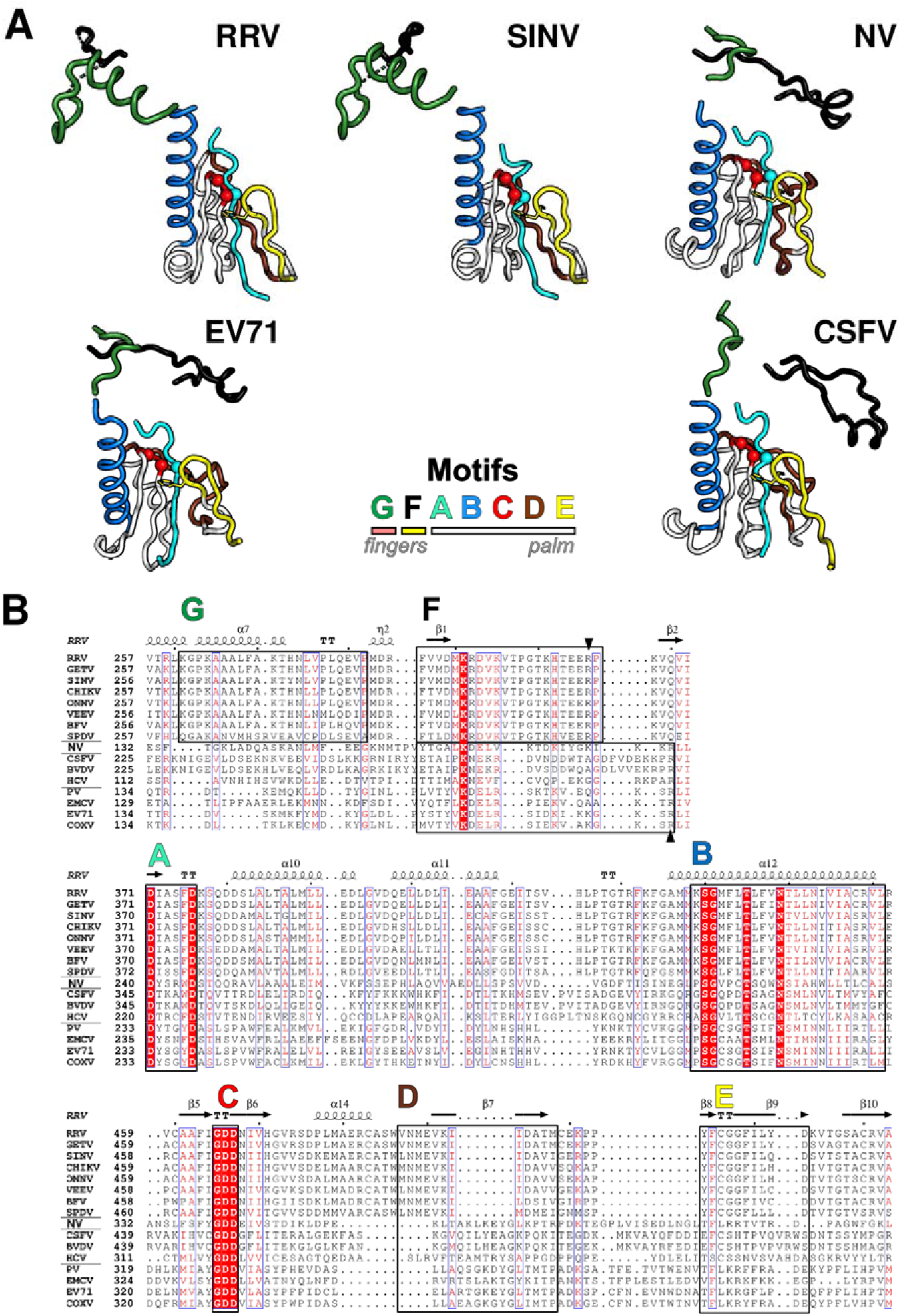
Comparison of the structure of alphavirus nsP4 with those of other viral RdRps. (A) The RRV RdRp^SER3^ (RRV, PDB 7F0S) and SINV RdRp (SINV, PDB 7VB4), and their shared top structural homologs, including Norwalk virus (NV, PDB 5TSN), Enterovirus-71 (EV71, PDB 6KWQ), and classical swine fever virus (CSFV, PDB 5Y6R). The motifs of these PDB structures were structurally aligned for comparison. The summarized motif and subdomain arrangement at the center are colored the same as Figure 1. The conserved aspartate residues in motifs C and A are displayed as peptide carbon backbone Cα spheres, while the conserved phenylalanine residue in motif E is shown in stick format. (B) The conservation of RdRps of +RNA virus based on primary sequence and secondary structure presented through Espript*. The sequence and secondary structure of RRV nsP4 (genus *Alphavirus*, family *Togaviridae*) were compared to those of different alphaviruses (RRV-to-SPDV) and members of the *Caliviridae* (NV)*, Flaviviridae* (CSFV-to-HCV), and *Picornaviridae* (PV-to-COXV) families. The secondary structure is labeled on top of the alignment and shows only α-helices (spring), β-strands (arrow), and β-turns (TT). Motifs A-G are boxed and colored according to (A). [*Espript coloring: The red-boxed columns are the most conserved, while the blue-boxed columns classify amino acid conservation based on their similar functional groups. *Espript label: *Togaviridae* - Ross River virus (RRV), Getah virus (GETV), Sindbis virus (SINV), chikungunya virus (CHIKV), o’nyong’nyong virus (ONNV), Venezuela equine encephalitis virus (VEEV), Barmah Forest virus (BFV), and salmon pancreas disease virus (SPDV); *Caliviridae* - Norwalk virus (NV); *Flaviviridae* - classical swine fever virus (CSFV), bovine viral diarrhea virus (BVDV), and hepatitis C virus (HCV); *Picornaviridae* - poliovirus (PV), encephalomyocarditis virus (EMCV), enterovirus-71 (EV71), and coxsackievirus (COXV).

### RRV RdRp displayed polymerase activity

A previously described fluorescent RNA-based assay (11) was optimized to improve robustness and sensitivity and used for the analysis of the RNA polymerase activities of the obtained recombinant proteins. In this primer extension assay, RNA intermediates (RIs) result from the incorporation of 1-7 Adenosine nucleotide residues (A residues), while the RNA product (RP) is the fully extended RNA obtained by incorporation of 8 A residues on the T1 RNA template (**Figure 4A**). Reactions were run for 24 hours (h) with time-point sampling at 0, 2, 4, 6, 8, 10, and 24 h.

**Figure 4.**
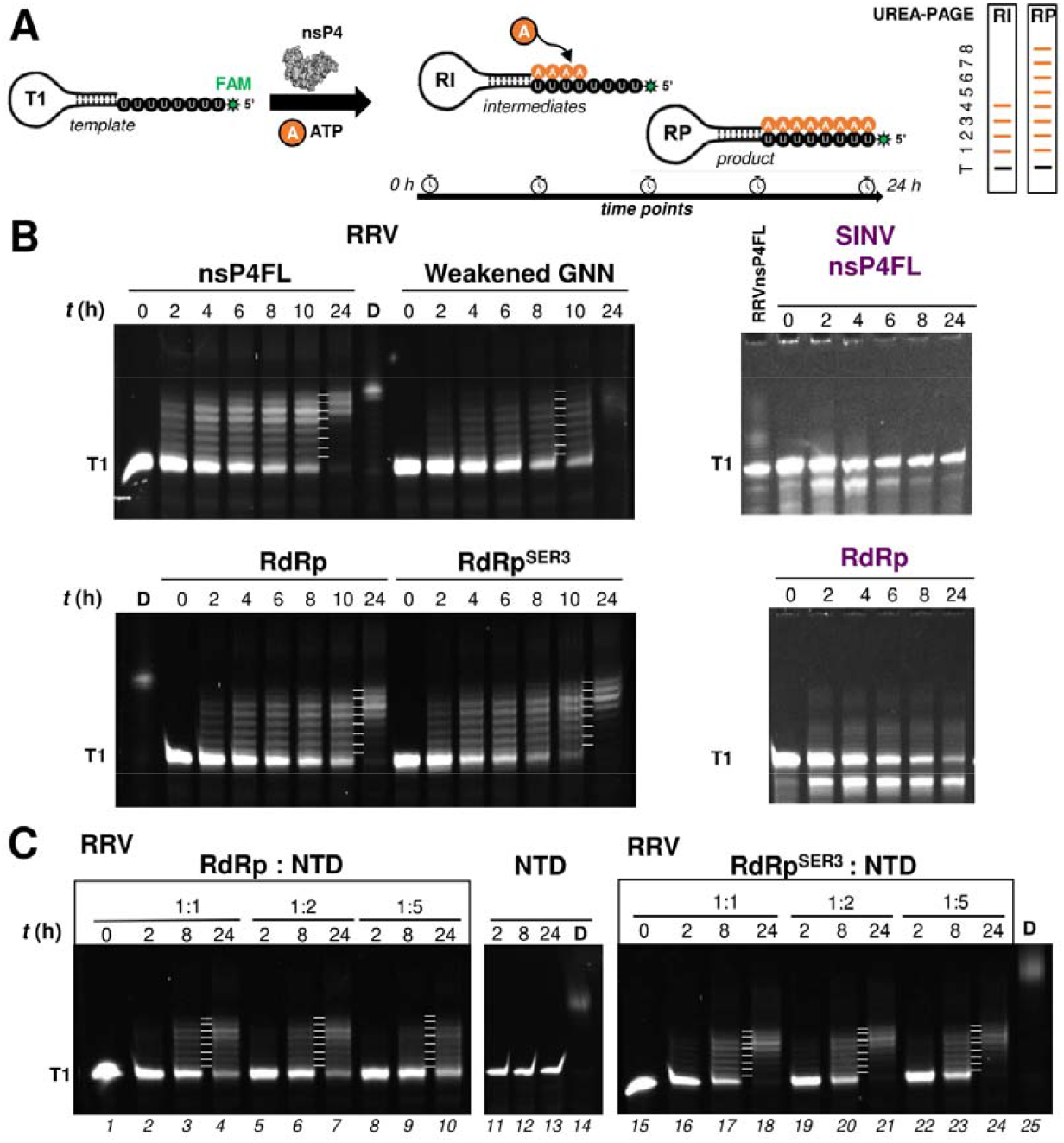
RNA polymerase activities of nsP4FL, RdRp, and RdRp^SER3^. (A) Schematic presentation of the discontinuous time point RNA polymerase reaction. Extension of fluorescently labeled (green star on the 5’ end) RNA template T1 by recombinant nsP4 protein (gray particle) results in ATP incorporations as A-residue (orange circle) into the RNA intermediates (RIs; a snapshot of one possible RI; right panel) and RNA product (RP; right panel), both of which are shown below with expected band migration with each band corresponding to the number of A-residue added to T1 on a UREA-PAGE gel. (B-C) The biochemical characterization of recombinant nsP4 proteins using discontinuous time-point gel-based polymerase assay was performed for 0-24 h. The denaturing UREA-PAGE gels are annotated with T1, time-point (t) in hour unit (hr), products made by dengue virus RdRp used as a positive control (named as D), and the tested recombinant nsP4 proteins from RRV (nsP4FL, polymerase weakened mutant of nsP4FL GNN, RdRp, RdRp^SER3^ and NTD) and SINV (nsPFL and RdRp, in purple text). For the polymerase reactions between RRV RdRp or RdRp^SER3^ and NTD, the stoichiometric molar ratio is shown in (C), that is marked with lane number for reference. The incorporation of each base is marked with a short white line shown between the last 2 time points.

The recombinant nsP4FL, RdRp, and RdRp mutants from RRV and SINV were enzymatically active and capable of extending the T1 template (**Figure 4 and S7**). The SINV nsP4FL and RdRp had weaker activities than those of RRV. The RIs were observed from 2 h onwards for all of these enzymes. The SINV nsP4FL is weaker than the RdRp protein probably caused by the high degradation rate of RNA template or RIs by the SINV nsP4 proteins (including their mutants) during the assay (**Figure 4B, right panel and Figure S7C-D**). Therefore, the following RNA polymerase assay and RNA binding characterization were focused on RRV nsP4 recombinant proteins that had minimal RNA degradation.

For all RRV nsP4 recombinant proteins, their polymerase reactions remained largely stagnant for over 10 h and subsequently accelerated to form a mixture of RP and RIs at 24 h (**Figure 4B, left panel and Figure S7**). After 24 h, nsP4FL ranks as the most efficient for the synthesis of RP and RIs with 6 or 7 A incorporations, followed by RdRp^SER3^ and RdRp. The conclusion that the activity of the surface-engineered mutant RdRp^SER3^ exceeded that of its wildtype (wt) counterpart was based on the observation that for the mutant enzyme, no RIs of less than 4 A added was detected at 24 h, while for its wild-type RdRp, a faint ladder consisting of the template and RIs with 1-3-A added was still visible even at this ending time point (**Figure 4B**). As expected, polymerase-weakened nsP4FL protein with GNN mutation at its motif C active site formed much smaller amounts of RIs, and the level of RP remained below the detection limit even at the 24 h time point (11,48,49).

To examine whether there is a functional interaction between the NTD and the RdRp domains, we measured the polymerase activity of nsP4FL and RdRp in the presence of the NTD protein at 1:1, 1:2, and 1:5 molar ratios. No primer extension was observed for the recombinant protein corresponding to the RRV nsP4 NTD (**Figure 4C: lane 11-14**). In the presence of the NTD protein, no increase in polymerase activity of RdRp^SER3^ or RdRp was observed at any time point (**Figure 4C**). Instead, an inhibitory effect of the NTD on the polymerase activity was observed more evidently for the RdRp (**Figure 4C: lane 4, 6, 10, 17, 20, 23**). Thus, the enhanced activity observed for nsP4FL could not be reproduced by mixing NTD and RdRp *in trans*.

### Activities of RRV replicase are affected by structure-guided mutations in nsP4

Having determined the crystal structure of RRV RdRp, we next analyzed the functional importance of several of its key structural features. As the introduction of unfavorable mutations into an infectious clone tends to result in no data (lethal phenotype) or the rapid appearance of revertants or compensating mutations (50,51), we instead used a novel two-component *trans*replication system (**Figure 5A**), which has extremely high sensitivity (25). Briefly, the system is based on the ability of separately expressed P123 and nsP4 of alphaviruses to form a functional RNA replicase. Coexpression of P123 and nsP4 with replication-competent template RNA also triggers the formation of RCs (spherules) similar to those formed in virus-infected cells (52), i.e., represents a relevant tool to study alphavirus RNA replication. Plasmids for the expression of RRV P123 and nsP4 as well as the corresponding template RNA have been described (25). The template RNA contains sequences coding for the following two reporters: firefly luciferase (Fluc) under the genomic promoter and *Gaussia* luciferases (Gluc) under the SG promoter (**Figure 5A**). For simplicity, the synthesis of full-length RNA serving as the template for Fluc expression is hereafter termed “replication”, and the synthesis of RNA serving as the template for Gluc expression is termed “transcription”. The efficiency of replication and transcription was estimated by the variations in corresponding reporter expression (53).

**Figure 5.**
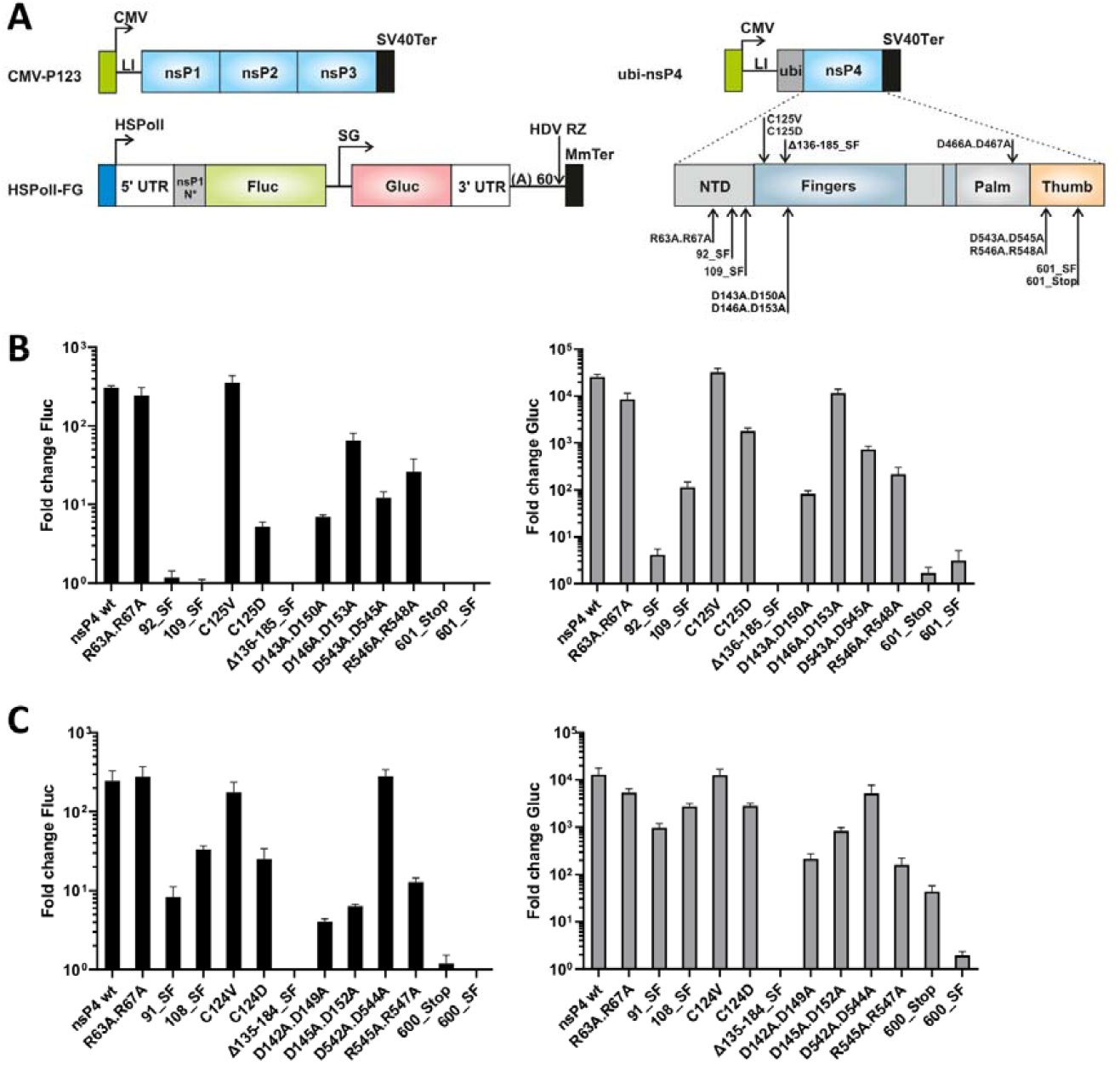
Effects of mutations in nsP4 on the activities of RRV and SINV *trans*-replicases. (A) The *trans*-replication system is composed of two plasmids (CMV-P123 and CMV-ubi-nsP4) for the expression of RRV and SINV replicase and a plasmid HSPolI-FG for the production of replication-competent RNA containing firefly luciferase (Fluc) and *Gaussia* luciferase (Gluc) reporters. In this assay, the increase in expression levels of Fluc and Gluc represented the production of full-length genomic RNA and SG RNA, respectively. HSPolI, a truncated promoter (residues −211 to −1) for human RNA polymerase I; 5’ UTR,full length 5’ UTR of an alphavirus; nsP1 N*, region encoding the N-terminal 77 (RRV) or 114 (SINV) amino acid residues of nsP1; SG, SG promoter spanning (with respect to termination codon of nsP4) from position −79 to the end of the intergenic region; 3’ UTR, truncated (last 110 residues) 3’ UTR of an alphavirus; HDV RZ, antisense strand ribozyme of hepatitis delta virus; MmTer, a terminator for RNA polymerase I in mice; CMV, an immediate early promoter of human cytomegalovirus; LI, leader sequence of the herpes simplex virus thymidine kinase gene with artificial intron; SV40Ter, simian virus 40 late polyadenylation region; Ubi, sequence encoding for human ubiquitin. Positions of mutations in nsP4 (residues numbered as in nsP4 of RRV) are indicated. (B, C) U2OS cells in 12-well plates were cotransfected with matching combinations of CMV-P123 and CMV-ubi-nsP4 or its mutant versions (in a 1:1 molar ratio) and the corresponding HSPolI-FG plasmids. As negative controls, CMV-ubi-nsP4^GAA^, which encodes nsP4 lacking RNA polymerase activity, was used instead of CMV-ubi-nsP4. Cells were incubated at 37°C and lysed 18 h posttransfection. Fluc and Gluc activities produced by wt and mutant replicases were normalized to the nsP4^GAA^ controls. Values obtained for nsP4^GAA^ controls were taken as 1. Means + standard deviation of three independent experiments are shown. (B) Effects of introduced mutations on RRV replicase activity. (C) Effects of introduced mutations on SINV replicase activity.

Mutations were introduced into nsP4 of RRV (**Figure S8**), and the activities of the replicates harboring mutant nsP4 proteins were compared with those of the wt replicase. Substitution of Arg63 and Arg67 with Ala residues did not result in significant changes in RNA replication, while transcription was slightly but significantly (p<0.001) reduced (**Figure 5B, Figure S8**). In contrast, only minimal transcription activity was detected for replicase harboring the Strep-FLAG tag after residue 92. The same tag inserted after residue 109 had a somewhat smaller impact; however, it still reduced the transcription level ~230-fold when compared with wt replicase (p<0.0001) (**Figure 5B**). Thus, the replacement of charged residues with Ala residues in the NTD of RRV nsP4 was well-tolerated compared to the insertion of peptide tags.

Replacement of Cys125 with the Val residue had no significant effect on the replication and transcription activities of the RRV RNA replicase. However, replacement of the same residue with Asp resulted in an ~60-fold decrease in replication and an ~14-fold decrease in transcription (p<0.0001 in both cases). Analysis of the nsP4 structure revealed that residues 136-185 of RRV nsP4 form a flexible loop. Replacement of this region with a Strep-FLAG tag resulted in the complete loss of RNA replicase activity, indicating a crucial function played by this region. This conclusion was corroborated by neutralizing the charges in this loop. Replacement of Asp143 and Asp150 with Ala resulted in a large ~43-fold decrease in replication and an even larger ~310-fold decrease in transcription (p<0.0001 in both cases). Interestingly, a similar replacement of Asp146 and Asp153 residues had a smaller impact: replication was only reduced ~5-fold (p<0.0001) (**Figure 5B**). Likewise, substituting charged amino acid residues with Ala in the thumb subdomain resulted in a decrease in RNA replicase activity: replacement of Asp543 and Asp545 caused an ~25-fold decrease in replication and an ~35-fold decrease in transcription (p<0.0001). The effect of substitution of the Arg546 and Arg548 residues caused a less prominent ~11-fold decrease in replication (p<0.0001), while the impact on transcription was more pronounced (~120-fold, p<0.0001) (**Figure 5B**). Finally, the introduction of a stop codon after residue 601, resulting in the loss of the ten C-terminal residues of nsP4 that could not be resolved in the present crystal structure, resulted in almost complete loss of RNA replicase activity. The insertion of a Strep-FLAG tag after residue 601 had a similar effect (**Figure 5B**). Taken together, these data indicate that the presence of the ten C-terminal residues and their location relative to the rest of nsP4 is crucial for RRV nsP4 activity.

## Discussion

The nsP4 polymerase crystal structures of two alphaviruses feature a conserved rigid fold in its palm subdomain but rather less so in the fingers and thumb subdomains (**Figure S3C**). This observation is unlikely due to crystal lattice packing forces as the two proteins were crystallized in different conditions and packed in different crystal lattices. The structures of alphavirus RdRps are likely the result of divergent evolution leading to the peculiar folds of the fingers and thumb at odds with the non-alphaviral RdRps. One of the most noticeable changes from both SINV and RRV RdRp crystal structures is the elongated β-strand adopted by motif D next to the catalytic GDD motif C located in the palm subdomain (**Figure 1 and Figure 4**). This structure contrasts with the canonical β-strand-loop motif found in other viral RdRps (**Figure 3A**). This elongated motif D with reduced flexibility could inhibit the NTP entry in the alphaviral RdRp apoenzyme state (42). Indeed, we observed no bound ATP or dATP but only the magnesium ions in the central catalytic cleft above the palm despite their presence in the crystallization buffer. More residues in the RRV nsP4 RdRp flex region are disordered than that of the SINV, likely due to the absence of the disulfide bond and crystal packing effect. Overall, structural analysis of the two RdRp structures suggests a partially unfolded, less active apoenzyme state exists for alphavirus nsP4 before RC assembly and viral RNA replication and transcription. It is expected that extensive structural rearrangement through the stages of the viral RNA biogenesis process, when nsP4 interacts with other nsPs and viral RNAs dynamically (42,54–57). The combination of HDX-MS, NMR, and Robetta homology modeling provided insights into the structural dynamics of nsP4 of both SINV and RRV in solution, especially for the NTD region. The homology model of RRV NTD features a mixture of disordered regions and a predicted helical secondary structure consisting of an HLH substructure (aa 53-74) (**Figure 2B-C**). Despite the challenging assignments of the cross-peaks to the NTD residues in the incomplete NMR spectra, these missing cross-peaks provided clues that the NTD may have active conformational exchanges (**Figure S6**). The HDX-protected region at the HLH substructure and the NMR-detected cross-peaks both suggest secondary structure elements in the NTD, which agree well with the NTD homology model. The HDX-MS analysis also indicates that the NTD does not interact with the catalytic RdRp domain of nsP4 (**Figure 2A**). The NTD of the viral RdRp is an adaptive domain that often undergoes divergent structural evolution to perform novel functions in various virus families or genera to target different hosts (54).

Analyses of RNA polymerase activity provide clues about the function of the nsP4 NTD. NTD is dispensable for viral RNA polymerase activity but appears to enhance *de novo* RNA synthesis for RRV nsP4FL (**Figure 4B**). The highest accumulation of fully extended RPs was observed when the NTD and RdRp domains were present *in cis* (i.e., in the context of the RRV nsP4FL). The NTD could also constitute an additional layer to regulate alphavirus RNA polymerase activity in a way reminiscent of how the fidelity of BVDV RdRp is modulated during RNA synthesis (54). Furthermore, the NTD from nsP4 could act as a chaperone required to first interact with nsP1-3 to mutually stabilize and organize into assemblies of active RC conformers to form the spherule for effective viral RNA synthesis. The SINV nsP4 was also previously shown in (23) to activate its viral RNA synthesis via interactions with nsP1-3. This may explain why the RRV and SINV nsP4 proteins alone showed weak and inefficient polymerase activity as compared to the positive control of dengue virus RdRp (**Figure 4, marked as D**). The disulfide bonding at C164-C323 in SINV nsP4 proteins which is suspected to cause misfolding of fingers and blocking the RNA binding tunnel could have further dampened the SINV polymerase activity with a greater degree of RNA degradation (**Figure 1**). The actively elongating RRV nsP4 recombinant proteins (**Figure 4**) are possibly the minor population, just like the CHIKV nsP4 previously reported in (11), that is stabilized with the help of the cations like manganese and magnesium and detergents like auryldimethylamine oxide (LDAO) and Triton X-100. In the absence of NTD, recombinant RdRp^SER3^ had slightly higher RNA polymerase activity than its wt counterpart, possibly due to its lowered surface entropy preventing precipitation to retain a higher amount of active protein.

To characterize the function of selected regions of the alphavirus RdRp for RNA replication and transcription in the cellular environment, a biologically relevant *trans*-replicase system was used (**Figure 5A**). The importance of the NTD for viral RNA replication was reflected by the strong negative impact caused by the Strep-tag inserted at various sites within the first 110 residues of nsP4. The flex region and the index fingertip are equally critical for viral RNA replication and transcription. We also found that several charged residues in the flex region (Asp143 and Asp150), in the fingers, and at the helix tip of the thumb (Arg543 and Arg545) are critical, presumably by providing electrostatic interactions either within the RdRp or/and with an RNA template. Finally, the last ten thumb residues protected from the solvent exchange in the HDX experiment were shown to be indispensable for RNA replicase activity (**Figure 5B-C**). These observations are consistent with the presence of an intramolecular interface within alphavirus nsP4 critical for viral RNA replication. In this respect, the C-terminus of the alphavirus RdRp could constitute a regulatory motif similar to that found in the flavivirus RdRp (42). Although nsP4 is the most conserved nsP, several known properties of recombinant nsP4 of SINV (12,23) indicate higher activity than what was observed for the RRV counterpart. We have also found that nsP4 of SINV and RRV is not fully functionally interchangeable: RRV P123 can form active RNA replicase with nsP4 either from RRV or SINV, while SINV P123 forms active replicase only with its cognate nsP4 (25). Based on sequence similarities, alphaviruses are divided into different complexes and clades (58). RRV is a member of the Semliki Forest virus complex, while SINV, the type member of the genus, belongs to the western equine encephalitis complex, making nsP4 of SINV a good candidate for validating to what extent findings made for nsP4 of RRV are applicable to nsP4 proteins of alphaviruses in general. Matching experiments performed using SINV *trans*-replicase provided results very similar to that of RRV (**Figure 5C**). The exceptions were mutations introduced in the NTD, which is the least conserved region of nsP4. The negative impact of the Strep-tag insertion was much smaller in the case of SINV nsP4: if the tag was introduced after residue 108, this insertion resulted in a relatively minor ~7-fold reduction in replication and a ~5-fold reduction in transcription. From a practical point of view, inserting an affinity tag at this position could be used for affinity pull-down of a functional SINV nsP4 and complexes formed by this protein. These findings are also consistent with our data, indicating that the NTD of nsP4 is flexible and not directly involved in RNA synthesis. In contrast, all mutations introduced into the RdRp domain of SINV nsP4 caused effects very similar to those observed for RRV. The largest difference between the two viruses was observed for the substitution of the Asp145 and Asp152 residues with Ala residues, which resulted in prominent reductions in both replication and transcription for SINV (by factors of ~38-fold and ~15-fold, respectively); these effects were significantly larger than those observed for corresponding mutations in RRV nsP4.

Our data indicate that there is structural and functional conservation in the RdRp domain even for distantly related alphaviruses. Therefore, it can be assumed that structural information for RRV and SINV RdRp is highly applicable for other alphaviruses and can be used for the rational design of drugs targeting many, or possibly all, medically important alphaviruses.

## Data Availability

The atomic coordinates and structure factors have been deposited with the Protein Data bank under accession codes 7F0S, 7VB4 and 7VW5.

## Supplementary Data

Supplementary Data are available at NAR online.

## Acknowledgment

We would like to thank the Protein Production Platform from NTU (PPP, NTU), NTU Institute of Structural Biology (NISB, NTU), Lee Kong Chian School of Medicine for providing the facilities and the general support from members of the D.L and A.M laboratories to complete our research. We thank the technical services provided by the “Synchrotron Radiation Protein Crystallography Facility of the National Core Facility Program for Biotechnology, Ministry of Science and Technology” and the “National Synchrotron Radiation Research Center”, a national user facility supported by the Ministry of Science and Technology of Taiwan, ROC – for data collection at Beamline NSRRC TPS 05A. We would also like to acknowledge the technical support from Japan SPring-8 Center RIKEN Targeted Proteins Beamline BL32XU and Australian Synchrotron, which is part of ANSTO, for the access to MX1/MX2 Beamlines (MX2 for the utilization of the Australian Cancer Research Foundation detector). Lastly, we thank the Paull Scherrer Institute (Villigen, Switzerland) for the provision of synchrotron beamtime at beamline X06DA of the SLS for data collection.

## Author Contributions

A. M. and D. L. designed research; Y. B. T., L. S. L., X. L., Y.-S. L., C. K., and J. Z. performed research; Y. B. T., L. S. L., X. L., Y.-S. L., C. K., J. L., J. Z., A. M., and D. L. analyzed data; and Y. B. T., L. S. L., A. M., and D. L. prepared the manuscript with input from all authors.

