## Supplemental file for "Crystal structures of alphavirus nonstructural protein 4 (nsP4) reveal an intrinsically dynamic RNA-dependent RNA polymerase fold"

**This document includes:**

Table S1

Figures S1 to S8

References for this SI

**Supplementary Table 1. Superimposition of the structures RRV and SINV RdRps.**

| | Residue Range* | | Residues considered | RMSD (super) |
|---|---|---|---|---|
| | **RRV** | **SINV** | **RRV/SINV** | |
| **Fingers** | 111-337, 409-431 | 104-336, 408-430 | 180/219 | 1.90 |
| **Palm** | 338-408, 432-513 | 337-407, 431-512 | 153/153 | 1.38 |
| **Thumb** | 514-600 | 513-601 | 77/89 | 2.05 |
| **Overall** | 111-600 | | 417 | 1.97 |

**Missing residues are not considered in the superimposition.*

**Supplementary figures.**

**Fig. S1. Anomalous Fourier map of the Seleno-Methionine (SeMet) residues on RdRpSER3.** The map is contoured at 3 σ, with each SeMet residue displayed as red sticks.

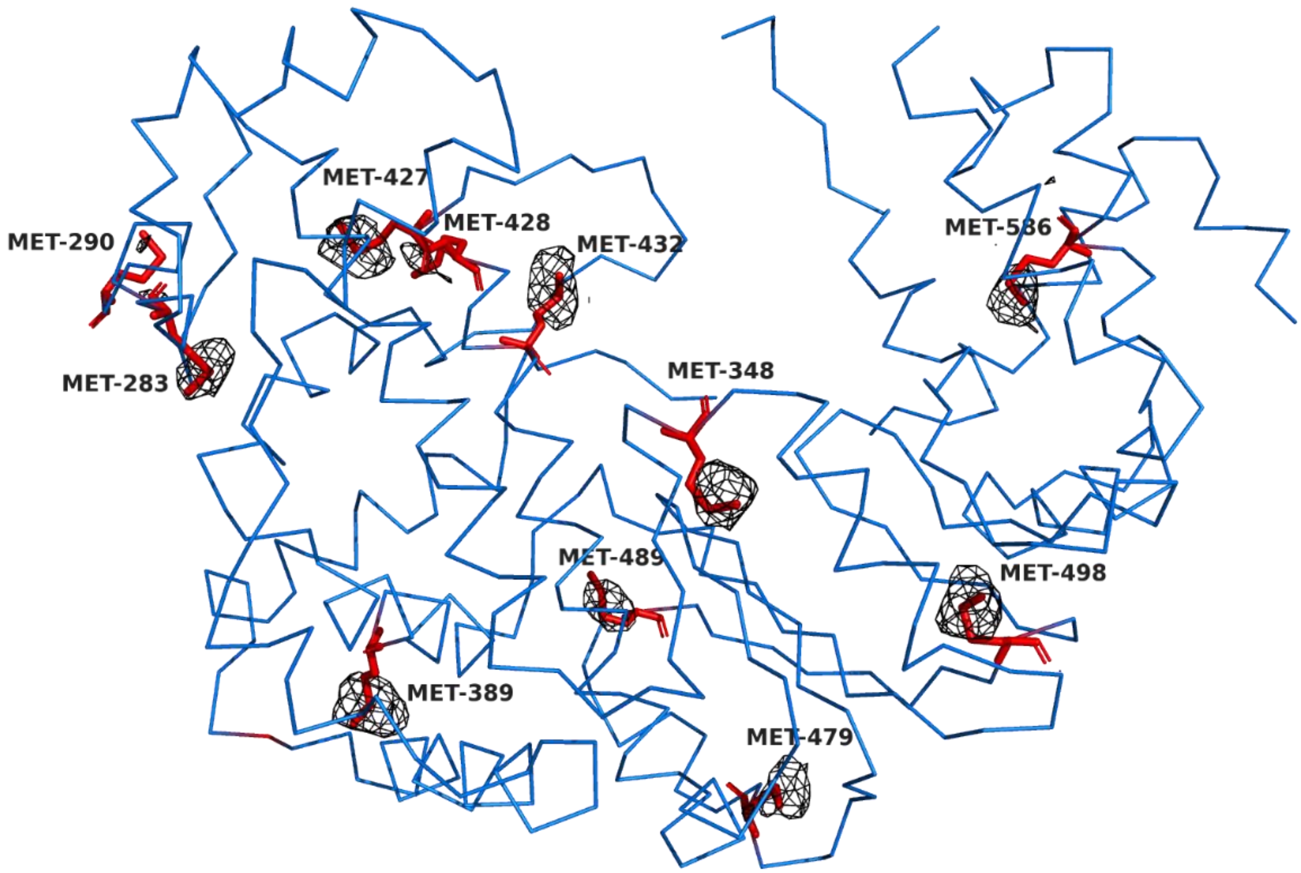

**Fig. S2.** The in-depth analysis of the built and disordered regions in the 2mFo-DFc map of RRV RdRp contoured at 1 σ, with reference to Figure 1. The blown-up views of the built model within the electron densities at (a) the ring finger, at (b) the thumb C-terminal region and at (c) the flex region (marked pink and built with polyalanine) which was hypothesized to insert itself through the central channel (teal arrow: insertion path labelled) like SINV RdRp. These maps portray the weak electron densities that discourage the building of residues. (d) The 2mFo-DFc map fitting was verified at the red motif C hairpin where GDD active site residues are. All missing residues were indicated next to each box.

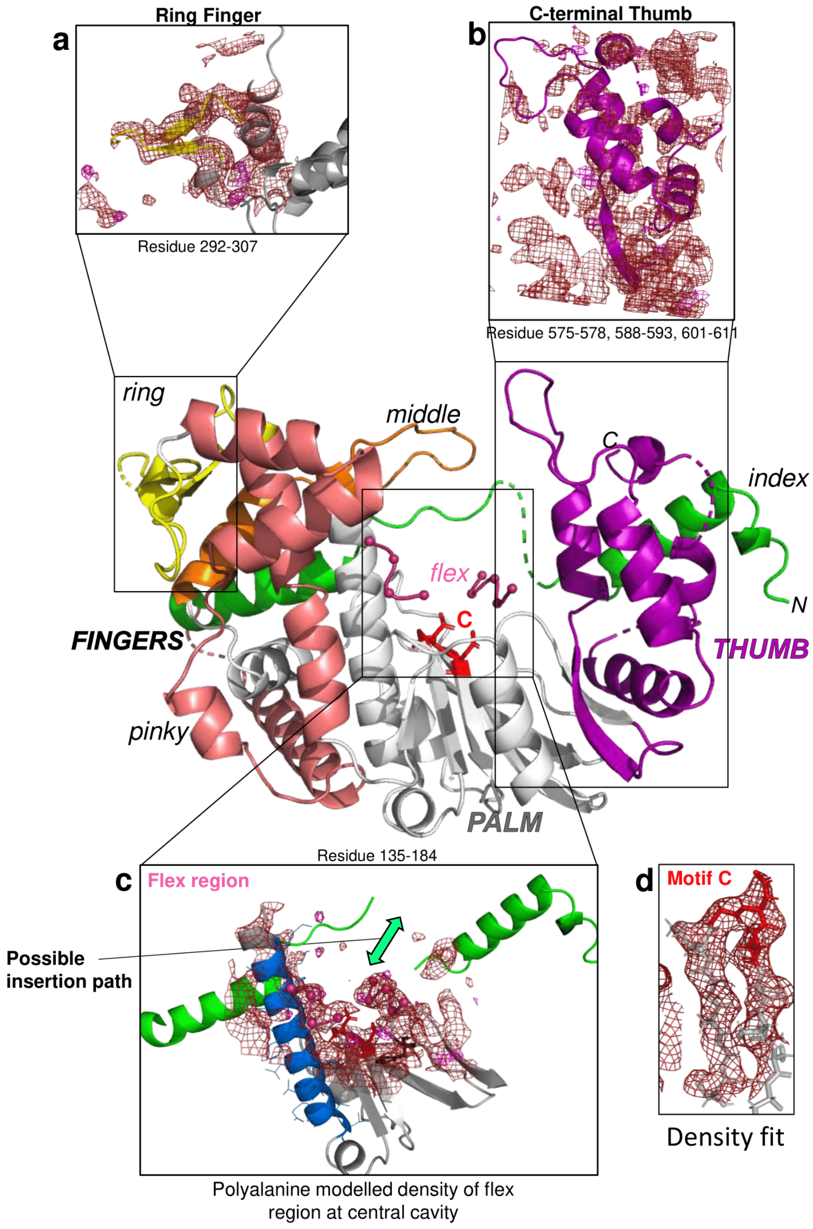

**Fig. S3.** The structural details of (A) two magnesium atoms (blue sphere) with each coordinated with four ordered water molecules (green sphere) and hydrogen bonding via carboxyl side chain of SINV structure aspartates (ASP392 and ASP466; oxygen atom in orange). The 2mFo-DFc maps were displayed at a contour of 1 σ. (B) Front view RdRp structure of monomeric RRV (gray) and (E) dimeric SINV (unit 1 in red and unit 2 in pink) are compared at the superimposed position as an overview of the interaction between index finger and thumb at different configurations. (C) Back view of the superimposed structures of RRV and SINV RdRps coloured as (B) with zoom in in-depth comparison for their structural differences at fingers (middle and index) and thumb.

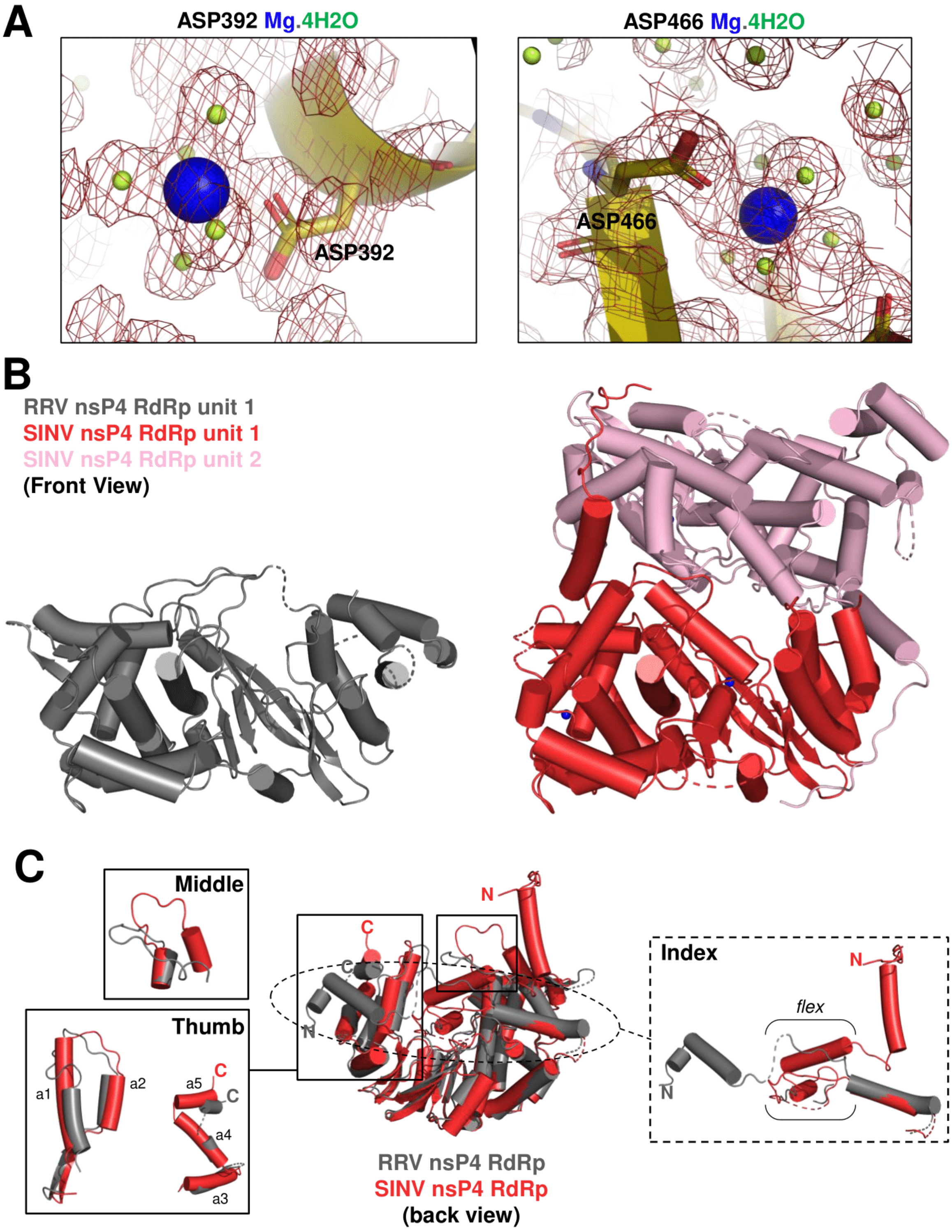
A
ASP392 Mg.4H2O
ASP392
ASP466 Mg.4H2O
ASP466
B
RRV nsP4 RdRp unit 1
SINV nsP4 RdRp unit 1
SINV nsP4 RdRp unit 2
(Front View)
C
Middle
Thumb
a1
a2
a5
C
C
a4
a3
N
C
C
N
RRV nsP4 RdRp
SINV nsP4 RdRp
(back view)
Index
N
flex
N

**Fig. S4.** HDX-MS 2D heatmap and heatmap-colored 3D models of RRV nsP4: (a) FL, (b) N (NTD), and (c) RdRp (NTD-lacking). The rainbow spectrum at the bottom of each panel represents the ascending percentage of deuteration-exchange (D%) from cool to warm colour.

**a**



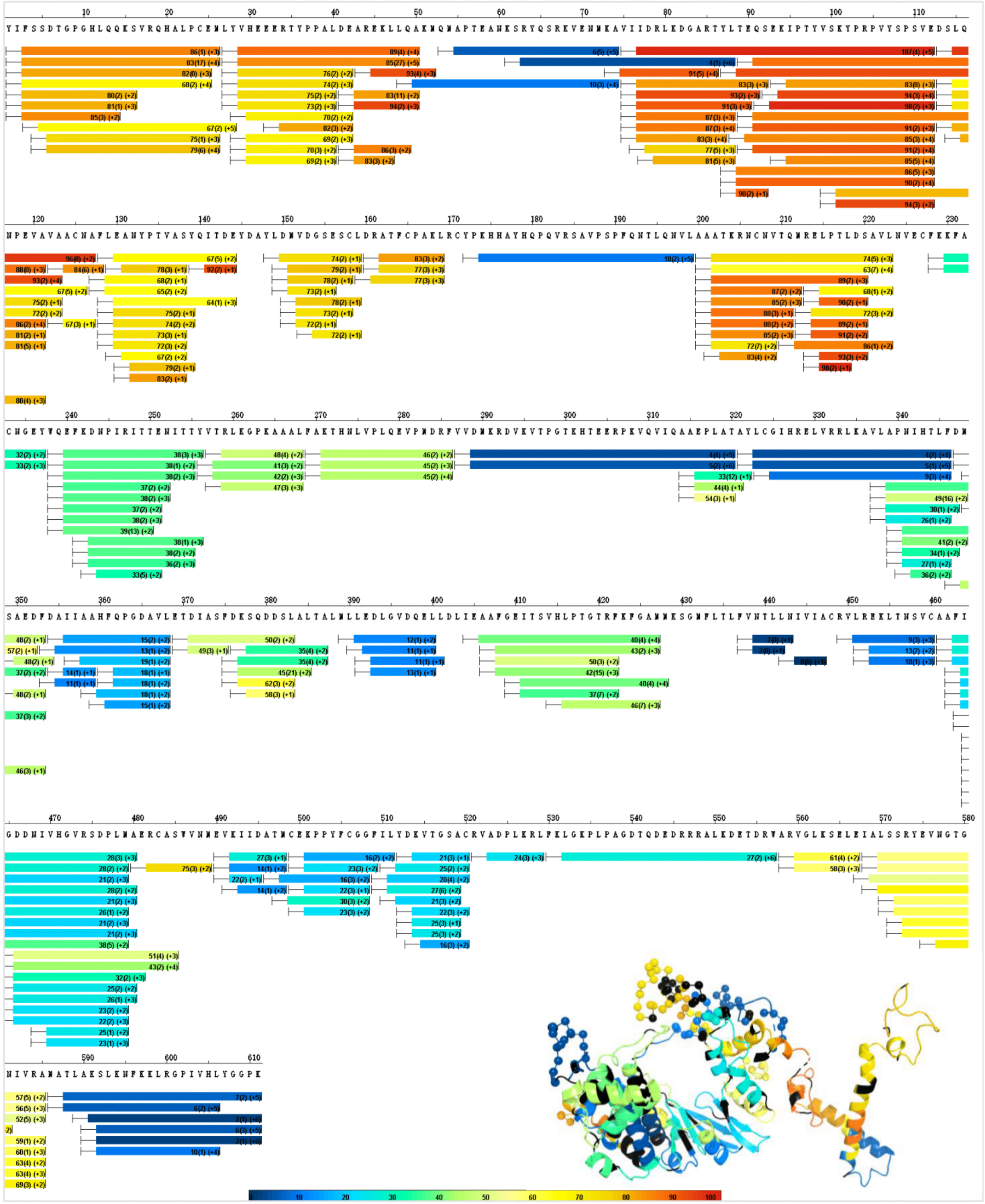

**b**

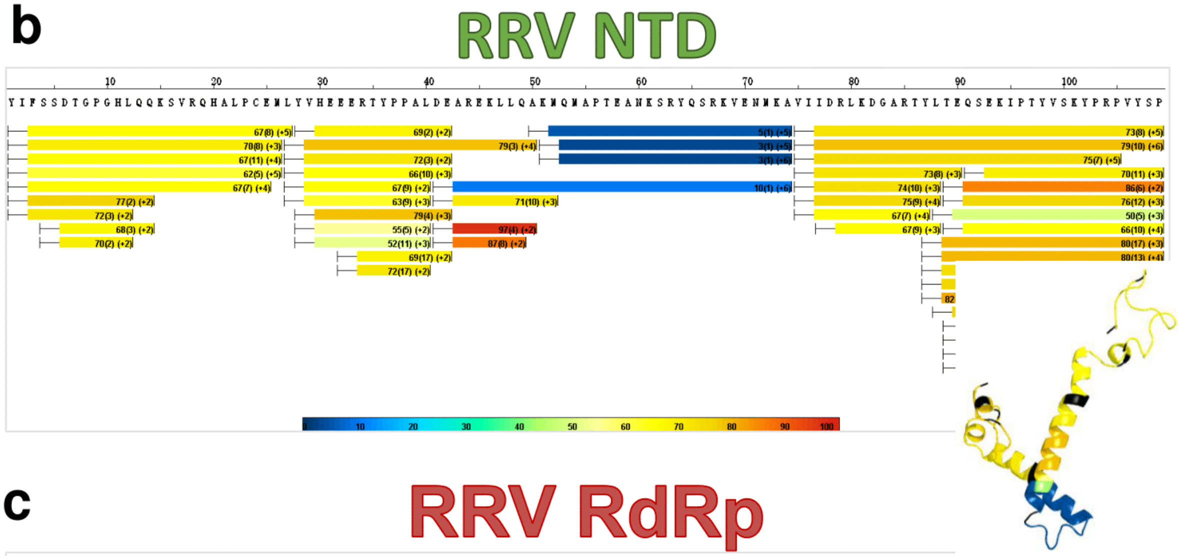
RRV NTD
10
20
30
40
50
60
70
80
90
100

**c**

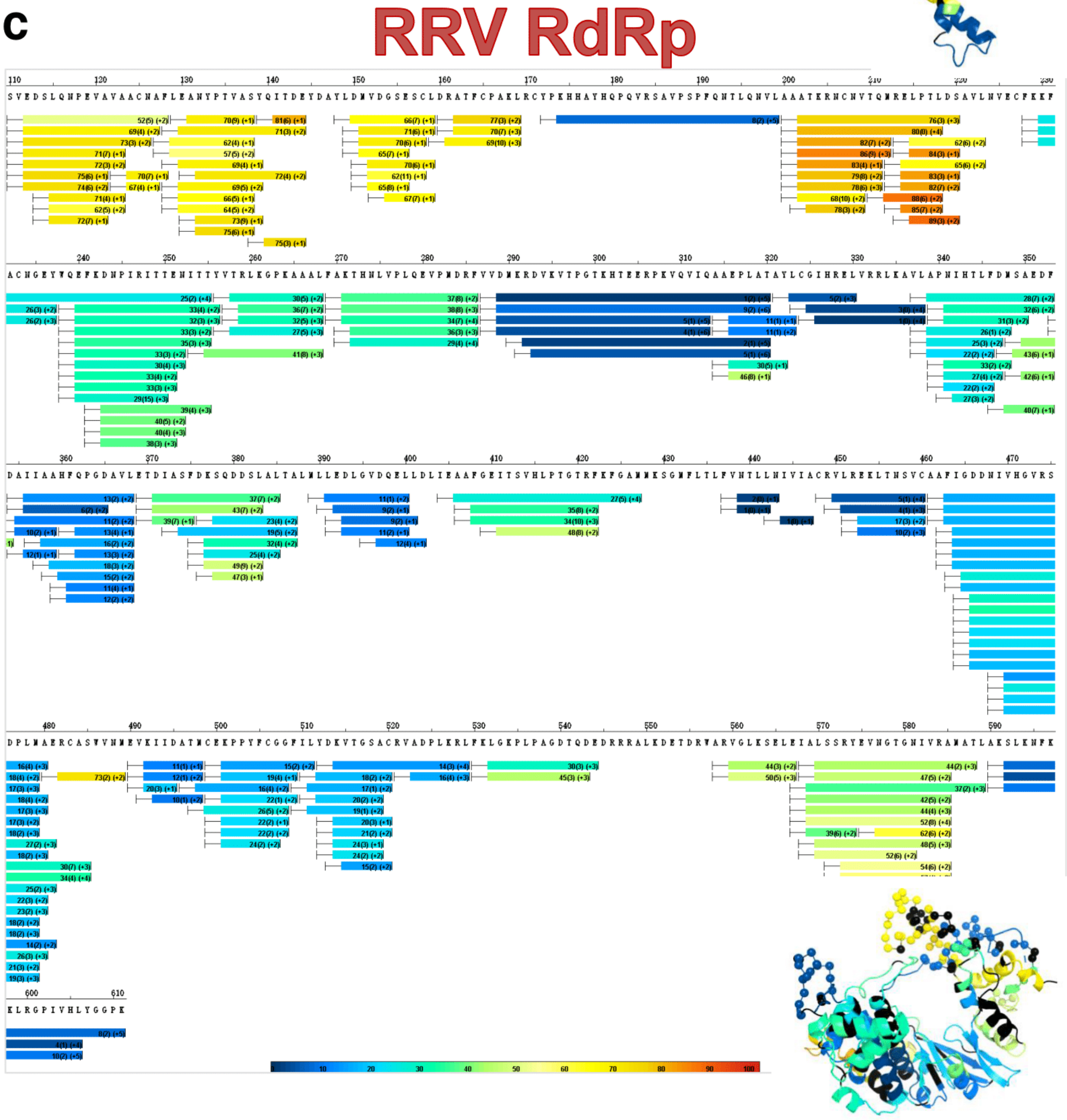
RRV RdRp
10
20
30
40
50
60
70
80
90
100

**Fig. S5.** HDX-MS 2D heatmap and heatmap-coloured 3D models of SINV nsP4: (a) FL, and (b) RdRp. The rainbow spectrum at the bottom of each panel represents the ascending percentage of deuteration exchange (D%) from cool to warm colour.

a

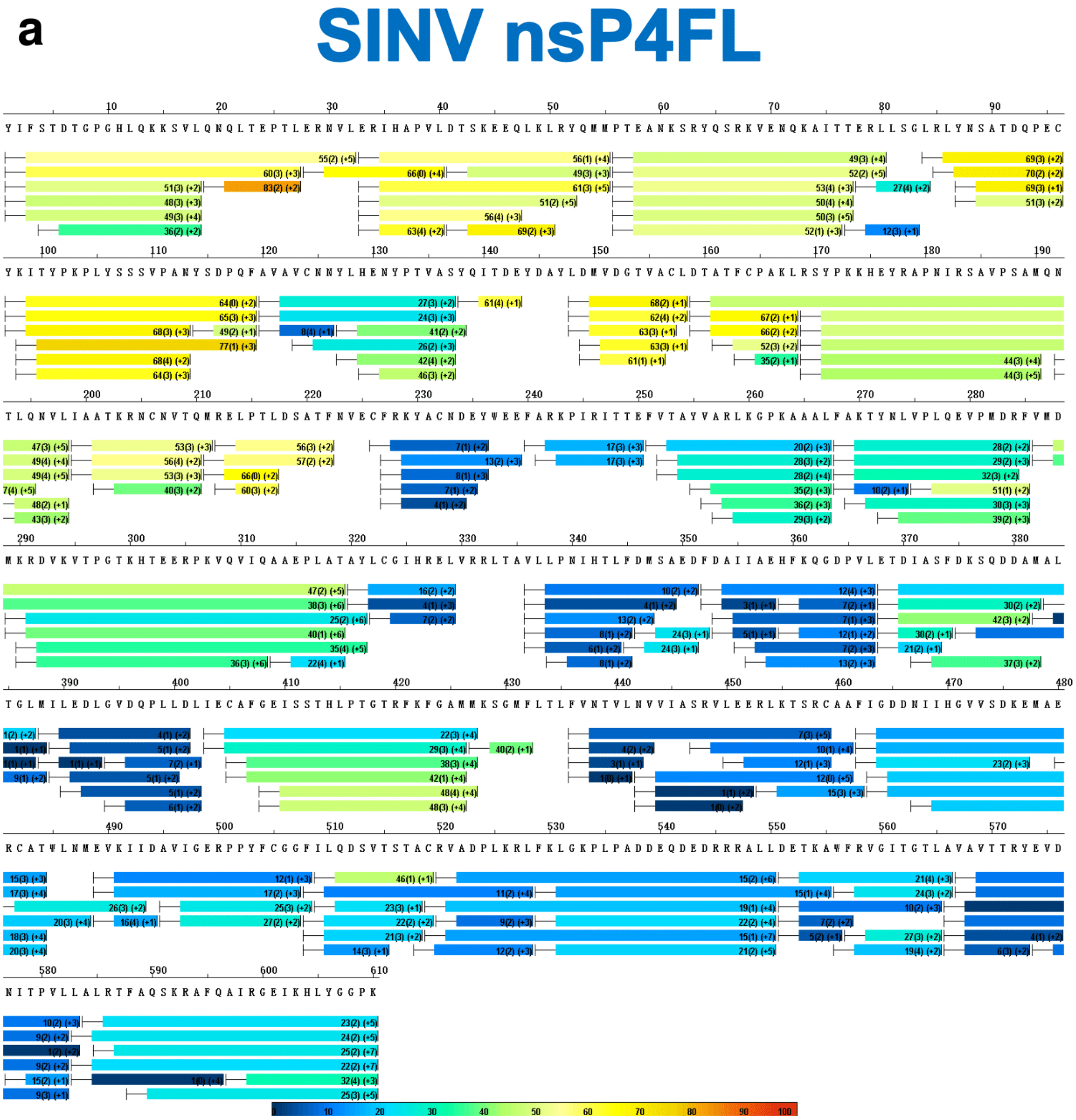


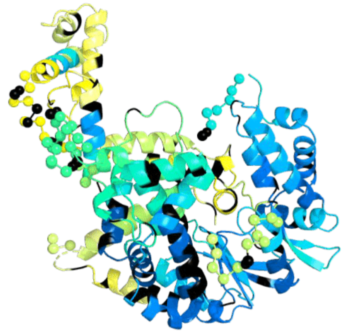

**b**

### SINV RdRp

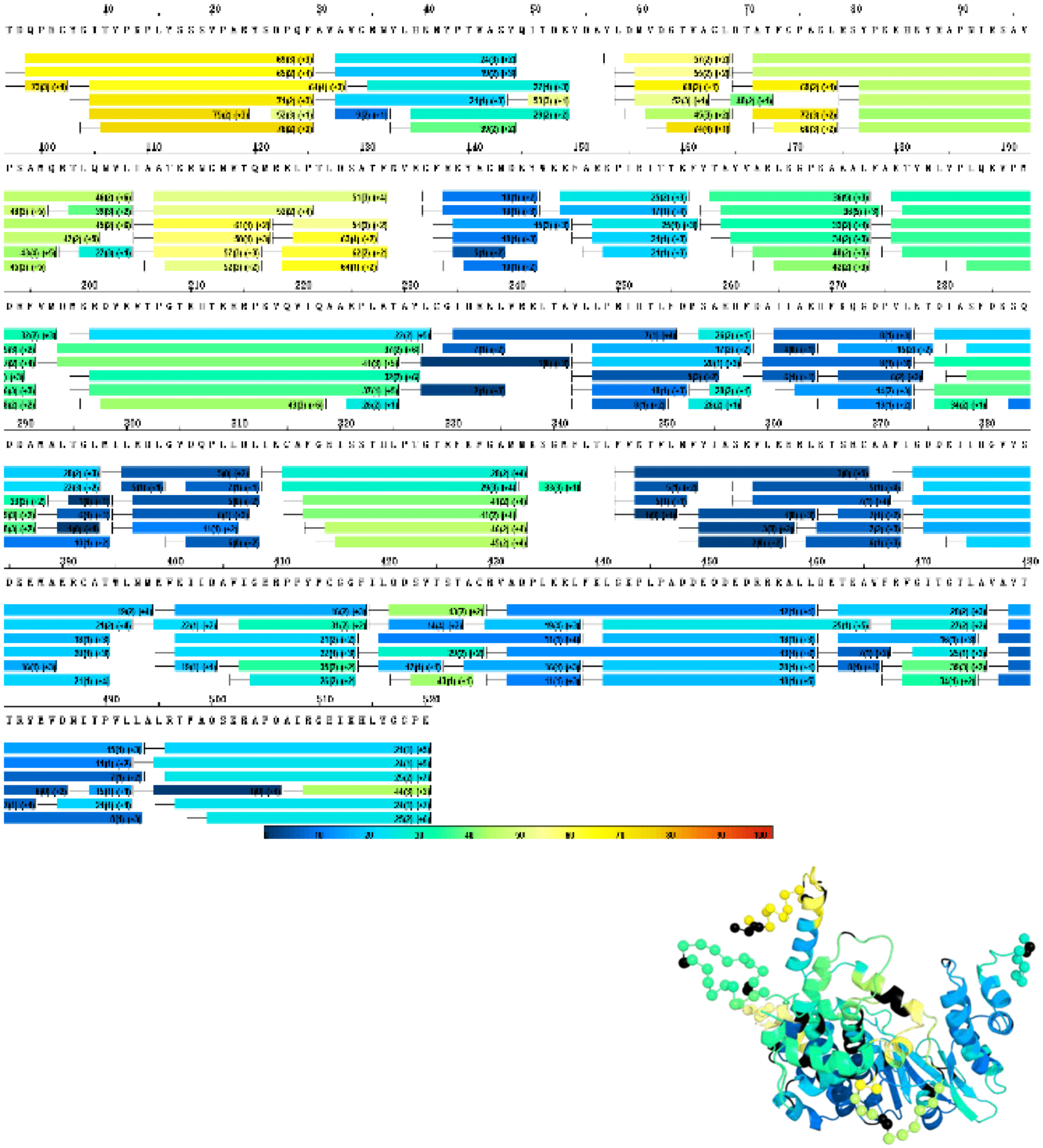

**Fig. S6**. $^{1}H$-$^{15}N$-HSQC spectrum of RRV NTD recombinant protein.

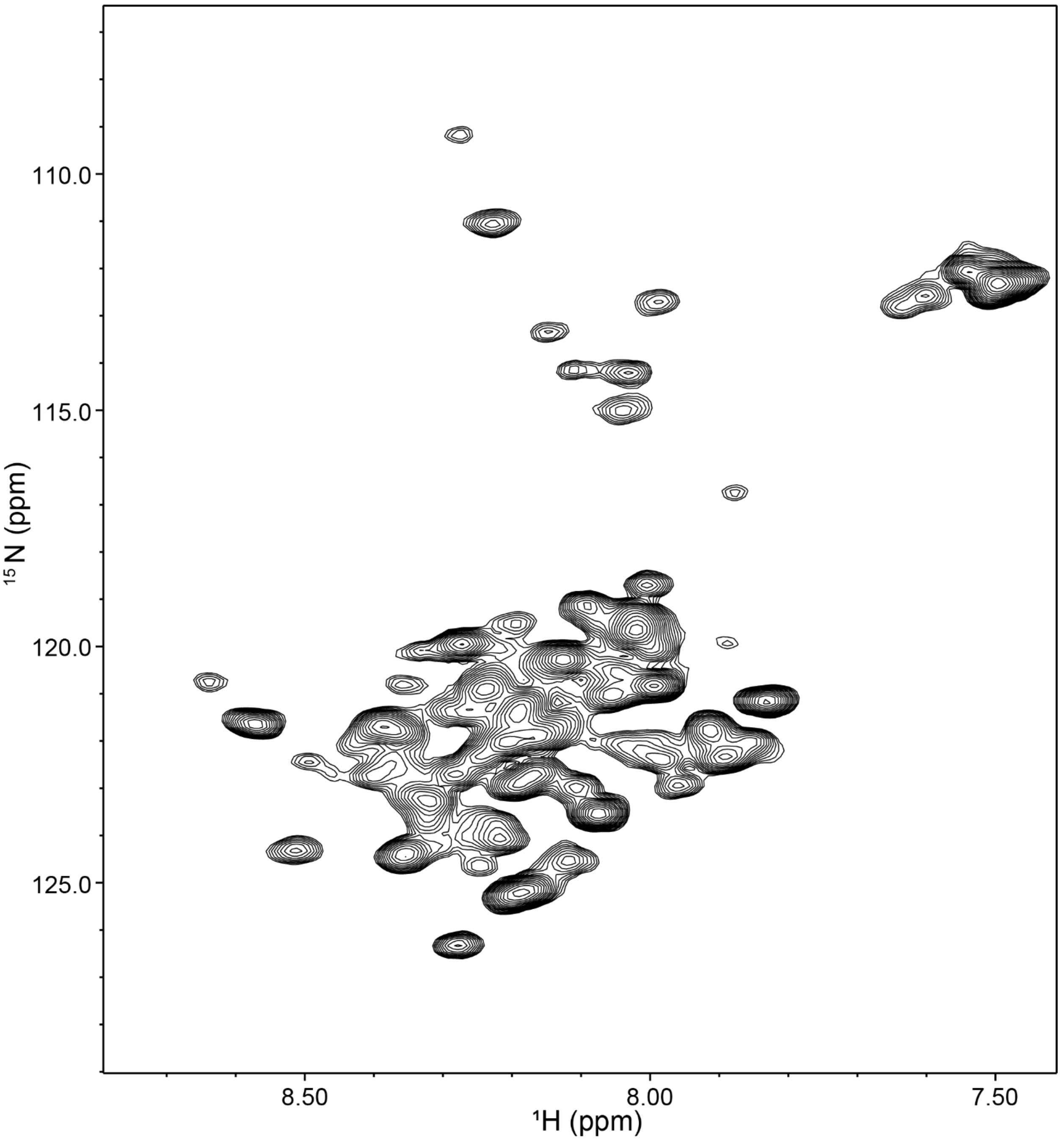

**Fig. S7**. RNA polymerase assay with full urea-PAGE gel image. The discontinuous-time-point polymerase assay represented in the uncropped gel images for different RRV recombinant proteins (a) nsP4FL, GNN, RdRp and $RdRp^{SER3}$, each sample from 0-24 h. (b) NTD (itself as negative control) and its complementary interactions with RdRp and $RdRp^{SER3}$ polymerases at different stoichiometric molar ratios (1:1, 1:2, 1:5) for 0-24 h. Dengue NS5 polymerase protein was used in the reaction as a reference (labelled as r). All reactions were based on RNA template T1. (c) The same polymerase assay experiment was repeated on SINV nsP4FL, RdRp and their C164 mutants: $RdRp^{CS}$ and $RdRp^{CN}$. (d) The multiple sequence alignment of alphaviruses shows the less conserved C164 among alphaviruses as a rational choice of mutation.

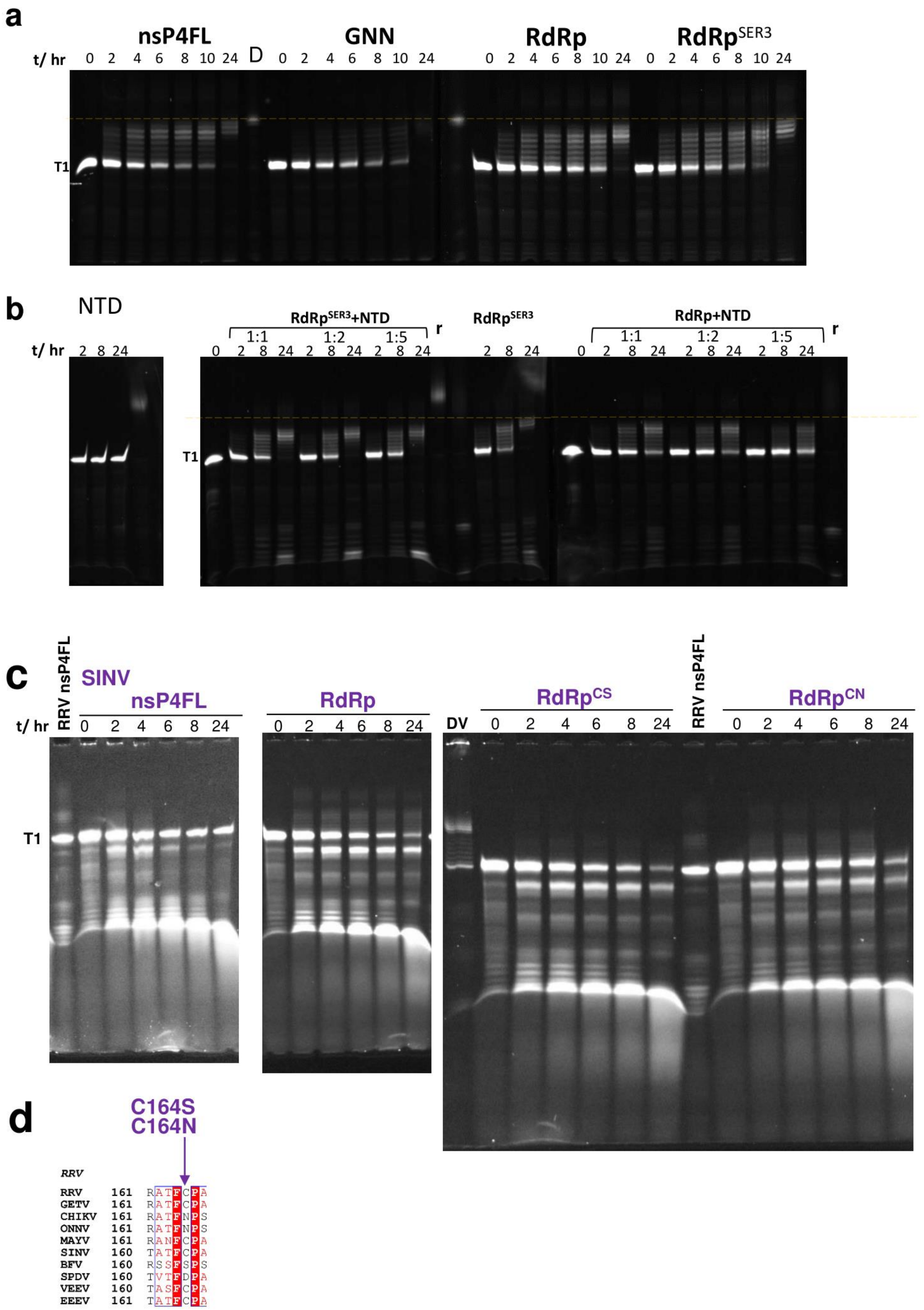

a
nsP4FL
GNN
RdRp
RdRpSER3
t/ hr
0 2 4 6 8 10 24
D
0 2 4 6 8 10 24
0 2 4 6 8 10 24
0 2 4 6 8 10 24
T1
b
NTD
t/ hr
2 8 24
RdRpSER3+NTD
1:1 1:2 1:5
r
0 2 8 24 2 8 24 2 8 24
RdRpSER3
2 8 24
RdRp+NTD
1:1 1:2 1:5
r
0 2 8 24 2 8 24 2 8 24
T1
c
RRV nsP4FL
SINV
nsP4FL
t/ hr
0 2 4 6 8 24
RdRp
0 2 4 6 8 24
DV
RdRpCS
0 2 4 6 8 24
RRV nsP4FL
RdRpCN
0 2 4 6 8 24
T1
d
C164S
C164N
RRV
RRV 161 RATFCPA
GETV 161 RATFCPA
CHIKV 161 RATFNPS
ONNV 161 RATFNPS
MAYV 161 RANFCPA
SINV 160 TATFCPA
BFV 160 RSSFSPS
SPDV 160 TVTFDPA
VEEV 160 TASFCPA
EEEV 161 TATFCPA

**Fig. S8.** The mutation sites are marked at **(a)** the alphaviruses'* sequence-conserved regions by targeting the regions not characterized by previous studies and with their design rationale and mutation positions tabulated in **(b)** for RRV and SINV. The secondary structure of RRV RdRp was presented on top of aligned sequences with α-helices (symbolized as α string for standard α-helix and η string for type $3_{10}$ α-helix) and β-strands (symbolized as arrow and TT for β-turns). **[OW: Old Word Alphavirus – Ross River virus (RRV), Getah virus (GETV), CHIKV, Onyong Onyong virus (ONNV), Mayaro virus (MAYV), Sindbis virus (SINV), Barmah Forest virus (BFV) and Salmon Pancreas Disease virus (SPDV); NW: New World Alphavirus – east equine encephalitis virus (EEEV) and Venezuela equine encephalitis virus (VEEV)]*

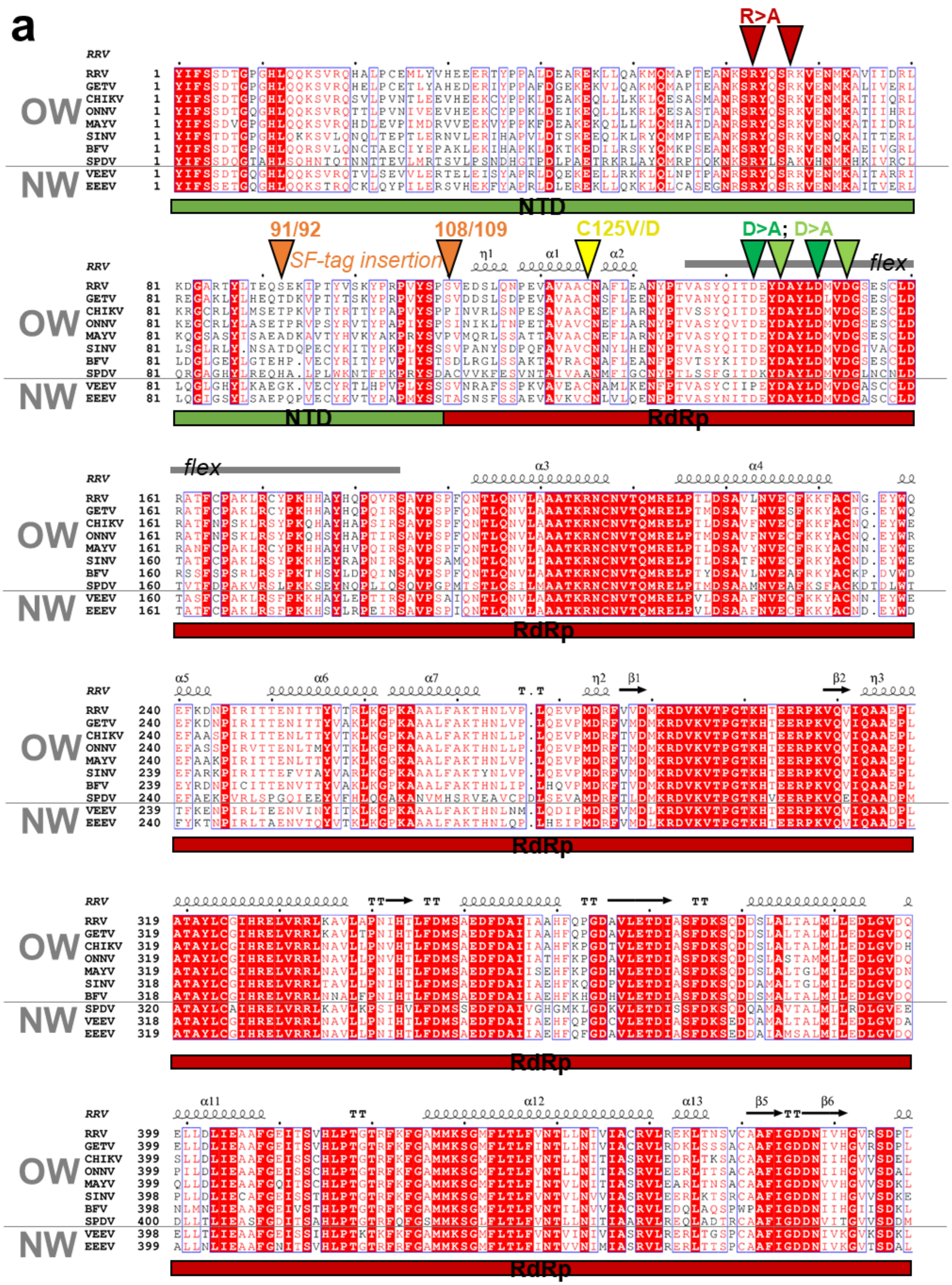
a
R>A
91/92
108/109
C125V/D
D>A; D>A
SF-tag insertion
flex
OW
NW
RRV
GETV
CHIKV
ONNV
MAYV
SINV
BFV
SPDV
VEEV
EEEV
NTD
RdRp

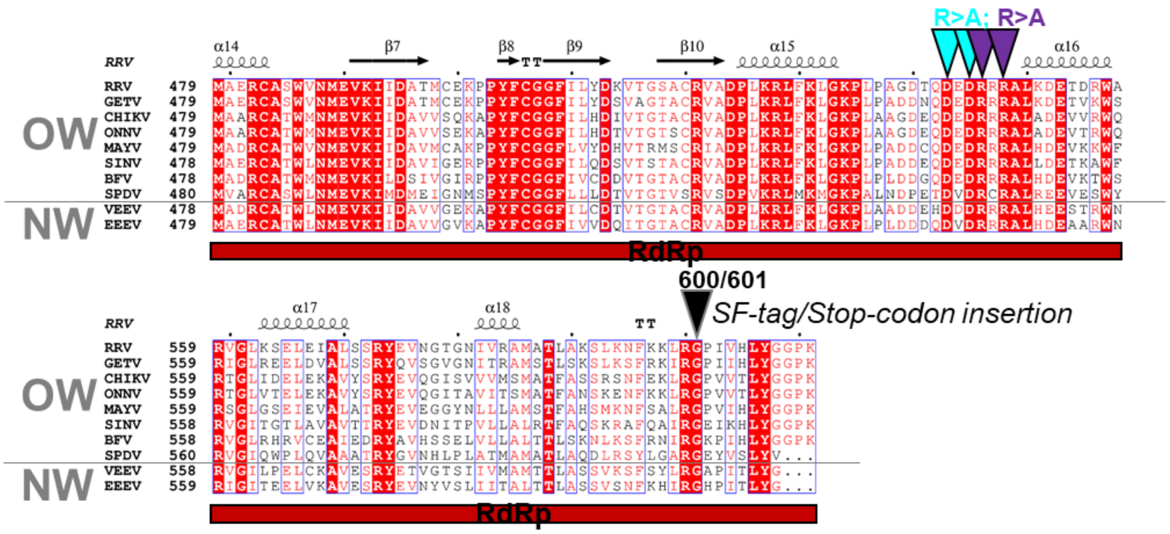


**b**

| Sub domain | Rationale | Type of change | RRV nsP4 (611 aa residues) | SINV nsP4 (610 aa residues) |
|---|---|---|---|---|
| NTD | HDX-protected area | Basic > Hydrophobic | R63A.R67A | R63A.R67A |
| | Tag insertion tolerance | SF-tag insertion | 92_SF | 91_SF |
| | Tag insertion tolerance | SF-tag insertion | 109_SF | 108_SF |
| Fingers | To probe Index-finger:Thumb Interaction | Cys > hydrophobic | C125V | C124V |
| | To probe Index-finger:Thumb Interaction | Cys > acidic | C125D | C124D |
| | Missing "flex" region – probe the tolerance to protein engineering and its possiblefunction | SF-tag insertion | Δ136-185_SF | Δ135-184_SF |
| | To challenge the highly conserved acidic-residue enriched region for their possible function in flex region | Acidic residues to alanine | D143A.D150A | D142A.D149A |
| | | Acidic residues to alanine | D146A.D153A | D145A.D152A |
| Thumb | Functional importance of charged helix tip at Thumb; could be contact to RNA | Acidic residues to alanine | D543A.D545A | D542A.D544A |
| | To probe function of charged helix tip at Thumb; could be contact to RNA | Basic residues to alanine | R546A.R548A | R545A. R547A |
| | To identify the structural and functional importance of missing C-terminus and tolerance to protein engineering | Truncation (stop codon insertion) | 601_Stop | 600_Stop |
| | | Strep-FLAG tag insertion after residue 601 (RRV)/600 (SINV) | 601_SF | 600_SF |
